# An oxo-mechanical coupling determines cell state

**DOI:** 10.64898/2026.03.12.711491

**Authors:** M Sreepadmanabh, Nivedita Hariharan, Dasaradhi Palakodeti, Tapomoy Bhattacharjee

## Abstract

Across myriad habitats and diverse lifeforms, oxygen and mechanics are the most ubiquitously varying and profoundly influential environmental regulators. While physiological niches feature both varying oxygen levels and heterogeneous mechanics, laboratory experiments typically interrogate these as independent variables. Here, we show that combinatorial regimes defined by varying oxygen partial pressures and environmental mechanics—an oxo-mechanical cue—induce functionally-distinct cellular states in 3D ECM-like contexts. Single-cell morphometrics combined with multi-omics reveal that cellular response to oxygen deprivation depends on external mechanical milieus, whereas, cellular engagement with different mechanical microenvironments depends on oxygen availability. Independently perturbing both hypoxic signaling and cytoskeletal activity further reveals a reciprocal oxo-mechanical regulatory coupling, which operates by differentially altering the global chromatin accessibility for transcriptional regulation in response to specific combinations of oxygen partial pressures and external mechanical milieus. Together, our findings establish that a coupling between oxygen and mechanics drives the emergence of microenvironmentally-defined cell states.

## 1. Introduction

The natural habitats of most lifeforms are complex 3D environments spanning simultaneously varying physical and chemical heterogeneities^1–6^. Consequently, cellular physiology and behavior are shaped by the dynamic integration of multiple extrinsic cues, rather than isolated stimuli^7–12^. However, current experimental paradigms typically perturb a single variable of interest while maintaining all else constant, which belie the complexity of combinatorial sensing encountered in vivo. Hence, despite extensive characterizations of the molecular effectors involved in environmental sensing and response, how mechano-chemically complex microenvironments modulate biological processes remains an open question. This gap is best exemplified by two of the most ubiquitous and broadly influential physical regulators – oxygen availability and environmental mechanics^13–18^(**Fig. 1a**). Considering that tissues across physiological and pathological scenarios invariably experience altered stiffness and restricted oxygen access, it is critical to decipher how such cues are interpreted at the cellular level (**Fig. 1b**)^15,19–43^. Oxygen profoundly impacts metabolism^14^, development^17,44,45^, differentiation^46^, and tumorigenesis^18^; while ECM mechanics influence motility^2,13,16^, morphogenesis^4,47^, fate transitions^48–50^, and metastasis^51^. However, almost all evidence for either oxygen or mechanics-driven regulation of living systems derives from experiments that independently perturb either of these two variables, without explicitly factoring in coordinate regulation^2,51–55^. Despite several instances of crosstalk between networks involved in hypoxic stress, ROS signalling, mechanosensation^56–59^, and ECM remodelling^60–62^, there exists no systematic understanding of how cells respond to concurrent variations in oxygen availabilities and ECM mechanics (**Fig. 1a**); and whether this combination of environmental cues merely act additively or antagonistically, or rather, give rise to an emergent class of unique cellular states – which cannot be inferred by interrogating oxygen and mechanics in isolation.

**Figure 1.**
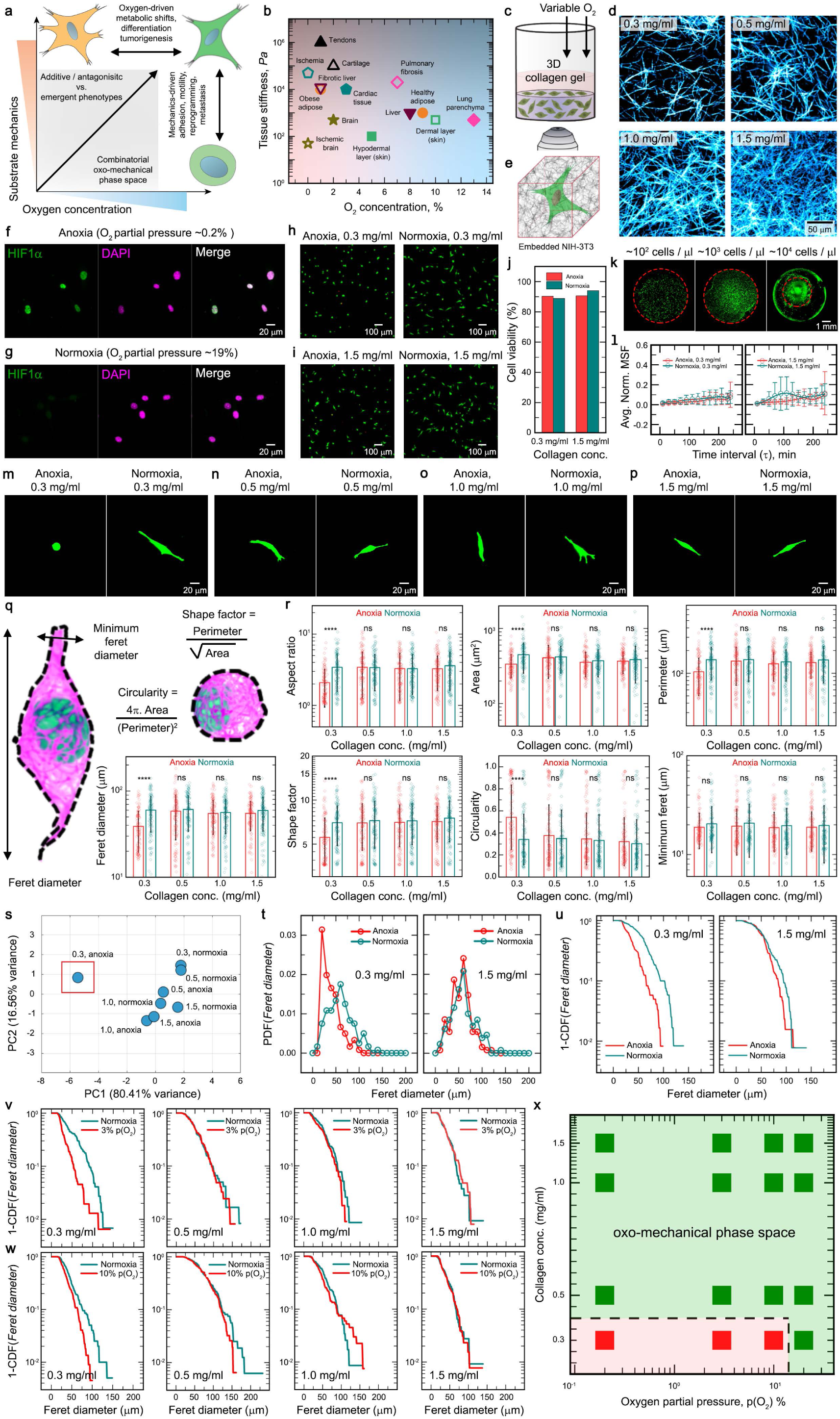
An experimental platform to interrogate cellular behavior across varying oxo-mechanical regimes reveals combinatorial control over morphology. (**a**) Present evidence documents how either of substrate mechanics or oxygen availability alters cellular behavior. However, the intersection of these physical components within the cellular microenvironment remains unexplored – i.e., an oxo-mechanical space. (**b**) Physiological niches typically exhibit spatiotemporally-varying material properties and oxygen levels, subjecting resident cells to diverse oxo-mechanical cues. (**c**) 3D culture platform for experimentally imposing oxo-mechanical regimes on NIH 3T3 fibroblast cells. Cells are homogeneously dispersed at low seeding densities within a 3D collagen type I hydrogel, which upon polymerization forms disordered fibrillar networks. Cell-laden hydrogels are subsequently incubated under different environmental oxygen partial pressures. (**d**) Collagen fibers imaged using reflectance microscopy show how the fiber density and porosity can be controlled by altering the relative fraction of collagen monomers in the precursor mix. (**e**) 3D collagen hydrogels offer a viscoelastic, mechanically tunable environment for probing cellular behavior across different oxo-mechanical regimes. (**f** and **g**) Immunostaining for HIF1α reveals that exposure to anoxia enhances nuclear localization of HIF1α, indicative of cellular hypoxic signaling responses. (**h**-**j**) A combination of anoxia and low fiber density collagen matrices do not detrimentally alter cell viability, which remains comparable across all four conditions tested herein. n > 70 individual cells per condition. (**k**) The low cell seeding densities used in this study (<100 cells / μL, where 1 μL = 1 mm^3^) ensure that cell-generated contractile forces do not mechanically deform the 3D collagen matrix. The region outlined in red indicates the size of the cell-laden collagen hydrogel following 24 hours of incubation under normoxic conditions. Only a 100-fold increase in cell density results in a significant contraction of the collagen hydrogel. (**l**) Representative time lapse micrographs and morphometrics performed on cells embedded within both poorly-reinforced (low fiber density) and well-reinforced (high fiber density) collagen matrices, and maintained under either anoxia or normoxia, following at least 12 hours of incubation under the respective conditions. No significant deviations from the initial morphology are observed, indicating that cellular conformations captured at least 12 hours post-experimental start time are stable snapshots. n >= 6 individual cells for all conditions, imaged using time lapse microscopy for = 4 hours each. Data represented as an average over all cells for the mean squared fractional change (MSF) in feret diameter for each cell between consecutive time frames, normalized to the feret diameter in the first frame of reference. (**m**-**p**) Representative maximum intensity projections of micrographs showing cellular morphology across different oxo-mechanical regimes. Barring a largely rounded-up population under anoxia coupled with the least mechanically reinforced (lowest fiber density) collagen matrices, cell morphology across all other tested conditions remains largely indistinguishable. (**q**) Schematic representation of the biophysical measurements extracted from maximum intensity projections of single cell micrographs. The feret diameter represents the longest dimension spanning the cell body, whereas, the minimum feret represents the shortest dimension spanning the cell body. (**r**) Quantifications for seven different morphometric parameters – aspect ratio, area, perimeter, shape factor, circularity, minimum feret, and feret – across eight different oxo-mechanical regimes, obtained by manually tracing the outlines of individual cells from maximum intensity projections of micrographs, with each condition including > 120 unique cells. All data shown here are sourced from samples prepared on the same day in order to minimize potential biological heterogeneity. The results clearly identify that only cells embedded within the least mechanically reinforced collagen matrix and maintained under anoxia exhibit quantifiably different morphometric profiles compared against cells from all other oxo-mechanical regimes. n >= 120 individual cells for each condition, data represented as mean +/− s.d. Statistical significance calculated using an unpaired t-test, with significance marked at a p-value < 0.0001. (**s**) Principal component analyses performed on a pooled dataset of all morphometric parameters across different oxo-mechanical conditions identifies the cellular population maintained under anoxia within the least-reinforced collagen matrices as a clear outlier. (**t** and **u**) Population-level comparison between cells cultured under both anoxia and normoxia within both the least- (0.3 mg/ml) and most- (1.5 mg/ml) reinforced collagen matrices, represented as (**t**) probability density functions and (**u**) as complementary cumulative distribution functions. Together, these analyses validate that only a combination of anoxia coupled with the least mechanically reinforced environment suffices to perturb the cellular morphology. In all other cases, additional reinforcement provided by the external mechanical milieu (by virtue of increasing fiber density) buffers the effects of anoxia. (**v** and **w**) Complementary cumulative distribution functions of cellular populations maintained under either (**v**) 3% or (**w**) 10% oxygen partial pressures, within different 3D collagen matrices, compared against control counterparts maintained under normoxic conditions. n > 100 individual cells for each condition. Barring the least-reinforced (0.3 mg/ml) collagen matrix, cells in all other mechanical regimes behave similarly across both normoxic and hypoxic conditions. (**x**) Combining morphometric analyses of the feret diameter across all the different oxygen partial pressures and 3D collagen matrices defines a phase space of cellular morphology across combinatorial oxo-mechanical regimes. This clearly demarcates a zone wherein low oxygen partial pressures coupled with the least mechanically reinforced collagen matrix together alter the cellular morphology. In all other cases, reinforced external mechanical milieus overcome the putatively detrimental effects of oxygen deprivation on the cellular morphology. Green squares represent similar morphologies between anoxic and normoxic samples; red squares represent a more rounded-up population morphology under anoxia as opposed to the more elongated population morphology under normoxia (p-value < 0.0001).

Here, by engineering mechanically tunable cell-laden 3D ECM-like collagen hydrogels and subjecting these constructs to varying oxygen availabilities, we study how distinct combinations of oxygen partial pressures and external mechanical milieus (an oxo-mechanical regime) alter cellular behavior. Combining single-cell morphometrics with transcriptomics and quantitative proteomics, we elucidate a phase space capturing differential cellular behavior across diverse oxo-mechanical regimes. Across these regimes, we discover a reciprocal oxo-mechanical coupling by independently modulating either the cytoskeletal state (via perturbation of actomyosin contractility and microtubular stability) or the hypoxic signalling (through tunable induction of chemical hypoxia). Functionally, this implies that cellular engagement with external mechanical milieus is determined by the hypoxic response, whereas, cellular response to varying oxygen partial pressures is determined by the intracellular mechanical state. The oxo-mechanical coupling is driven by globally-altered chromatin conformations, which manifest differentially across specific combinations of oxygen levels and ECM mechanics – resulting in an environmentally-responsive control over the magnitude and outcomes of transcriptional factor activity. This leads to a profound shift in cellular states, as captured by an integrated gene regulatory network model which identifies lynchpin molecular effectors driving oxo-mechanical regulation. Together, we demonstrate that cellular states are an emergent property of environmental oxo-mechanical regimes.

## 2. Results

### 2.1. Oxygen partial pressures and ECM mechanics combinatorially regulate cellular morphology

To experimentally recapitulate the complex and disordered architecture of ECM-rich 3D cellular microenvironments, we employ collagen type I hydrogels (**Fig. 1c**-**1d**). Within these, NIH-3T3 fibroblasts are homogeneously dispersed (prior to polymerization) at low seeding densities, generating cell-laden 3D matrices (**Fig. 1e**). To engineer controlled oxo-mechanical regimes, we simultaneously vary both the mechanical properties of the collagen hydrogel and the environmental oxygen partial pressures. By varying the monomer concentration between 0.3 mg/ml and 1.5 mg/ml, we modulate both the viscoelastic properties as well as the fibre density of the 3D matrix (**Fig. 1d** and **SI Fig. 1a**). Further, to span a range of environmental oxygen levels, we employ sealed temperature-controlled (37°C) culture chambers, within which the oxygen partial pressures are varied between anoxic (∼0-0.2% oxygen), hypoxic (∼3% to ∼10% oxygen), and normoxic (∼19-20% oxygen) conditions (**Fig. 1f-1g**). Cell-laden collagen hydrogels supplemented with nutrient media are incubated within these environmentally-controlled chambers, effectively enabling an interrogation of cellular behavior across a range of oxo-mechanical regimes. We confirm that these culture conditions do not detrimentally affect the cellular viability over experimental time periods of up to 24 hours (**Fig. 1h-1j**). During this interval, it is critical to conserve the culture conditions. While oxygen partial pressures can be held constant, collagen hydrogels can be collectively deformed by cell-generated contractile forces, leading to abruptly altered cell density and modification of material properties. We avoid this by employing cellular densities which are ∼1/100 of the thresholds required to mechanically deform the collagen hydrogel matrices (**Fig. 1k**). Together, these data support the suitability of this experimental strategy as a tunable, controlled, and stable method of subjecting cells to varying oxo-mechanical environmental regimes.

Next, we systematically assess cellular morphology across several oxo-mechanical regimes. First, using time lapse imaging and quantitative morphometrics, we verify that observed morphologies are not a transiently varying phenotype, but rather, stable representations of the cellular behavior under specific oxo-mechanical regimes (**Fig. 1l** and **SI Fig. 1b**). Typically, adherent NIH-3T3 fibroblasts assume a spindle-like morphology in collagen hydrogels. While most of the oxo-mechanical regimes demonstrate this pattern, interestingly, we observe an alteration of the morphology occurring exclusively within the lowest fibre density (0.3 mg/ml collagen) anoxic conditions wherein we find predominantly rounded-up cells (**Fig. 1m-1p** and **SI Video 1-4**). However, our prior measurements of viability confirm that this morphological shift is not due to apoptosis. To quantitatively describe these effects, we separately obtain z-stack images using confocal microscopy for hundreds of individual cells across each oxo-mechanical combination. We use maximum intensity projections of these z-stack micrographs and manually trace the outlines of each such cell to quantitatively characterize several morphometric attributes describing its shape (**Fig. 1q-1r** and **SI Fig. 2**). Given the heterogeneity in cell morphologies, we consciously avoid automated image processing algorithms in order to maximize the accuracy of structure segmentation. Further, considering that systemic variability introduced by single-cell heterogeneity may render ensemble-averaged statistics fallacious, we verify the robustness of our sampling technique. Our analyses reveal that for the oxo-mechanical regimes under consideration, a sample size of approximately 60 cells is truly representative of the population behavior (**SI Fig. 3**). Considering this, we maintain a minimum sample size of 100 cells for all morphometric analyses. Additionally, principal component analyses (PCA) of the data pooled together from the multi-parametric morphometrics for anoxic and normoxic cells maintained across different mechanical regimes also validate these findings (**Fig. 1s** and **SI Fig. 4a**), as well as help identify the feret diameter – the longest dimension spanning the cell body – as a direct and sufficient representation of cellular morphology across the tested conditions.

**Figure 2.**
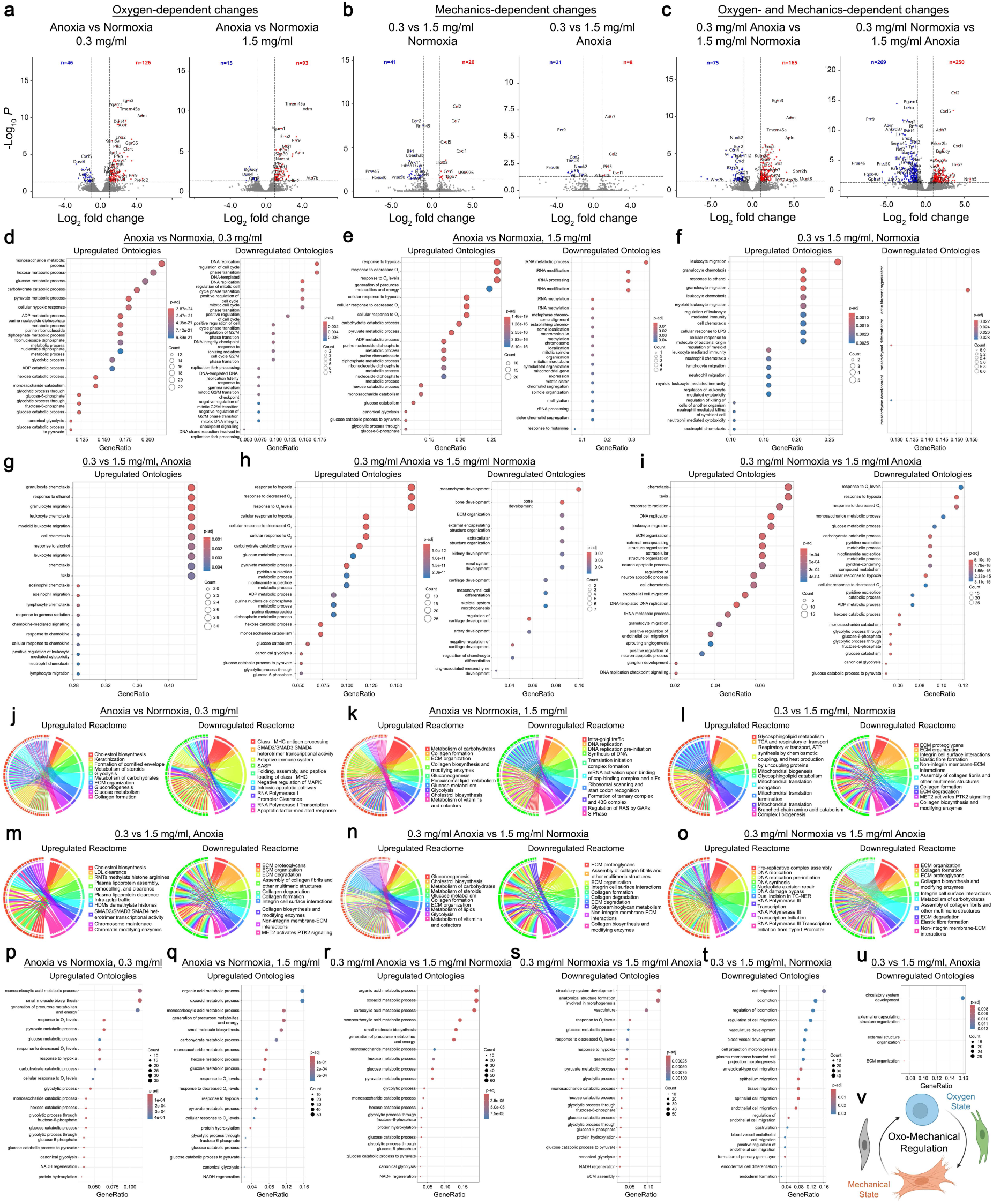
A reciprocal oxo-mechanical coupling determines cellular states. (**a**-**c**) Volcano plots from transcriptomic analyses comparing cells maintained across different combinations of oxygen partial pressures and environmental mechanics – demonstrating that oxo-mechanical regimes induce fundamentally different cellular states, characterized by distinctly different, microenvironment-dependent gene expression programs. (**d**-**i**) Ontological associations with biological processes (either upregulated or downregulated) based on differentially-expressed genes across variations in either oxygen partial pressures, environmental mechanics, or both variables together. Statistical thresholds for identifying differentially-expressed genes across all pairwise comparisons were set as |log2FC| >= 1, i.e., 2-fold change in expression levels, with adjusted p-value < 0.05. (**j**-**o**) Reactome analyses based on quantitative proteomics, highlighting differentially-enriched pathways (either upregulated or downregulated) across various oxo-mechanical combinations. (**p**-**u**) Ontological associations with biological processes (either upregulated or downregulated) based on differentially-abundant proteins across variations in either oxygen partial pressures, environmental mechanics, or both variables together. Statistical threshold for identifying differentially-abundant proteins across all pairwise comparisons was set at a p-value < 0.05. No additional fold-change cutoff was imposed here, on account of the comparatively lower detection range for proteomics as opposed to the RNA-Seq above – hence, to aid the explorative and descriptive nature of these analyses, we have included all differentially abundant proteins retained after imposing the p < 0.05 statistical testing cutoff. (**v**) A reciprocal oxo-mechanical regulatory coupling governing the cellular state. The coupling links the intracellular mechanical state and the intracellular oxygen signaling state as reciprocally-interacting coupled modules, the interaction between which determines the overall cellular state across different environmental oxo-mechanical regimes.

**Figure 3.**
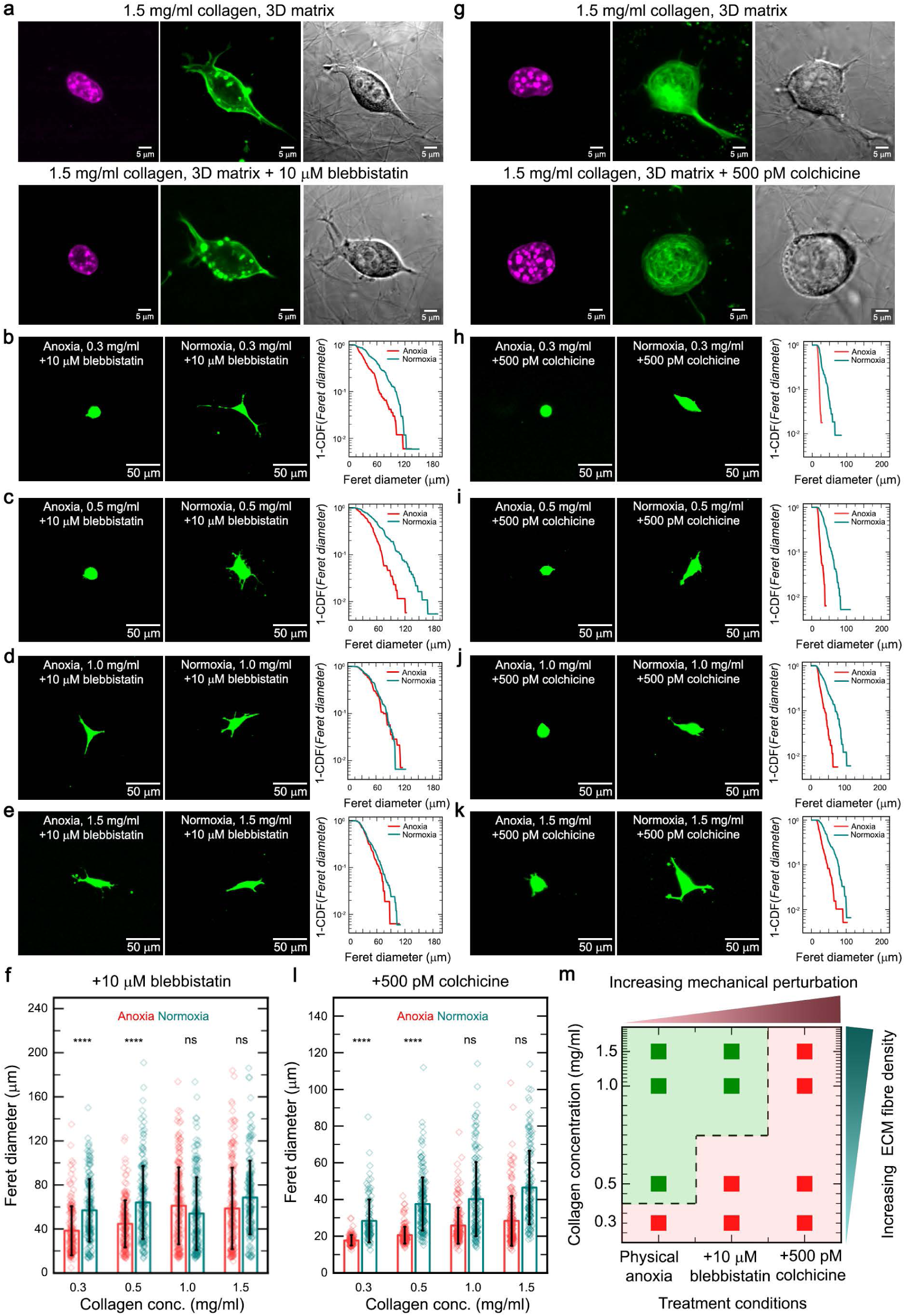
The intracellular mechanical state determines cellular response to environmental oxygen availability. (**a**) Blebbistatin treatment (10 μM) alters actomyosin contractility without significantly altering the cortical actin organization for cells adherent in 3D collagen hydrogels. (**b**-**f**) Effect of perturbed actomyosin contractility on cellular morphology induced by blebbistatin treatments across different oxo-mechanical regimes. Mechanically perturbed cells under these conditions show a difference between the anoxic and normoxic conditions in the 0.5 mg/ml collagen matrix as well, whereas previously these differences remained restricted to only the least-reinforced (lowest fiber density, 0.3 mg/ml) collagen matrix. n > 160 individual cells for each condition, data in bar plots represented as mean +/− s.d. Statistical significance calculated using an unpaired t-test, with significance marked at a p-value < 0.0001 (**g**) Colchicine treatment at picomolar concentrations (500 pM) minimally, but not entirely, disrupts microtubule organization for cells adherent in 3D collagen hydrogels. (**h**-**l**) Effect of perturbed microtubule stability induced by colchicine treatments across different oxo-mechanical regimes, showing a significant shift in cellular morphology across all external mechanical regimes under anoxic conditions. n > 100 individual cells for each condition, data in bar plots represented as mean +/− s.d. Statistical significance calculated using an unpaired t-test, with significance marked at a p-value < 0.0001. (**m**) Phase space of cellular morphology for different intracellular mechanical states across different collagen fiber density matrices, compared between anoxic and normoxic samples, both subject to the same chemical treatments where specified. Green squares represent similar morphologies between anoxic and normoxic samples; red squares represent a more rounded-up population morphology under anoxia as opposed to the more elongated population morphology under normoxia (p-value < 0.0001). Three distinct conditions are represented here – unperturbed mechanical state under only physical anoxia, perturbed actomyosin contractility under blebbistatin treatment coupled with physical anoxia, and perturbed microtubule stability under colchicine treatment coupled with physical anoxia. The phase boundary shifts across these three contexts prove that the intracellular mechanical state directly determines the cell’s response to different oxygen availabilities, thereby validating one half of the proposed reciprocal oxo-mechanical coupling.

**Figure 4.**
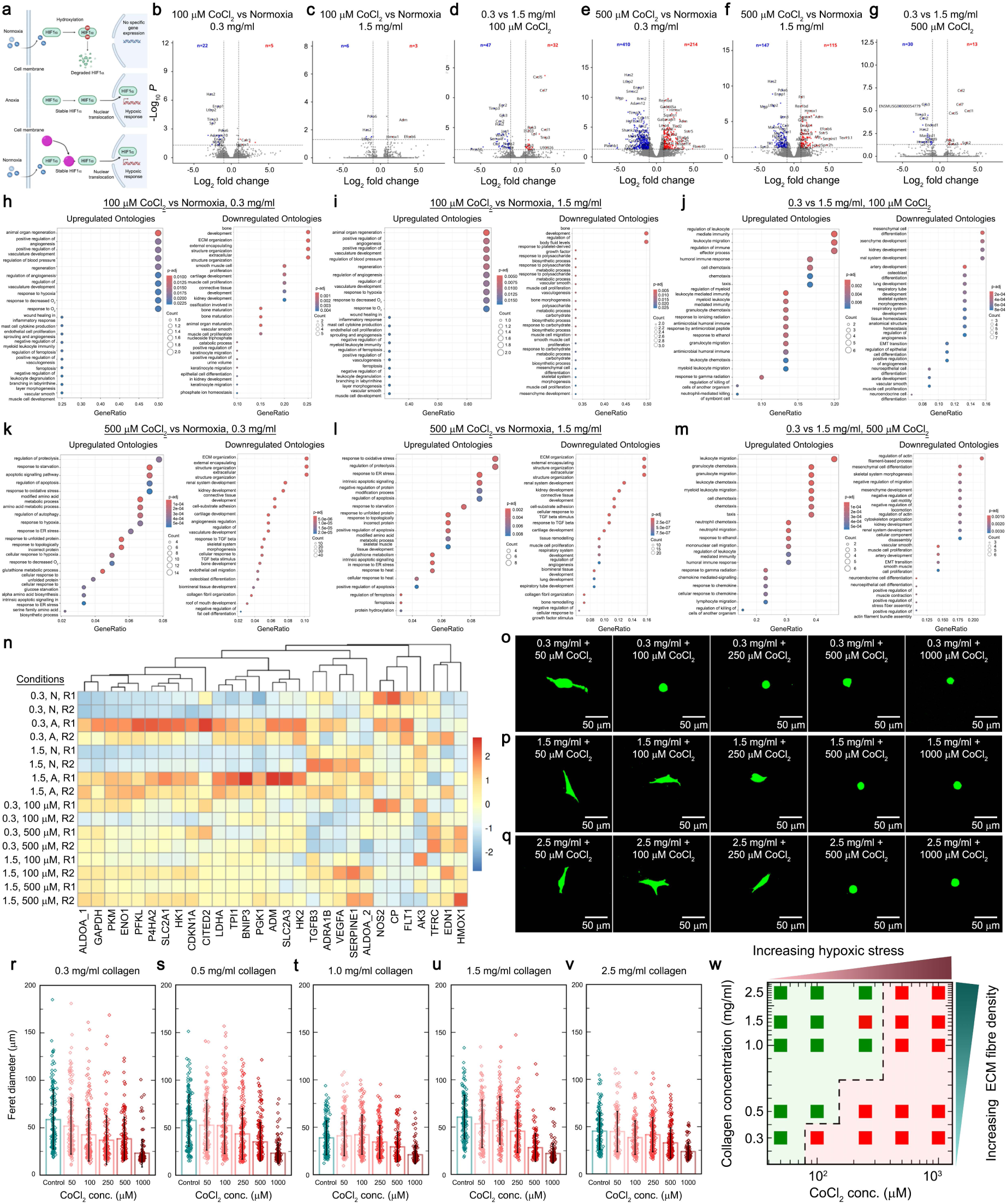
The intracellular hypoxic signaling determines cellular engagement with different external mechanical regimes. (**a**) Schematic representation of low oxygen partial pressure sensing via the canonical hypoxia-induced factor (HIF1α). Presence of molecular oxygen leads to hydroxylation of the HIF1α protein, targeting it for degradation. Un-hydroxylated HIF1α translocates to the nucleus, where it acts as a broadly-active transcription factor, triggering genes involved in cellular hypoxic responses. Cobalt chloride acts as a chemical hypoxia mimic by stabilizing the HIF1α molecule even in the presence of molecular oxygen, allowing a dosage-dependent control over hypoxia-induced gene expression without necessitating an alteration to the environmental oxygen levels. (**b-g**) Volcano plots from transcriptomic analyses for two different grades of cobalt chloride treatments – 100 μM and 500 μM – reveal that cellular states are combinatorially regulated by both microenvironmental mechanics and chemically-induced hypoxic responses. The increase in number of differentially expressed genes going from 100 μM to 500 μM shows that chemically-induced hypoxic stress is also dosage-dependent. The change in differential gene expression appears most pronounced for cells embedded in poorly-reinforced mechanical milieus while being exposed to a high degree of chemical hypoxia, consistent with the findings using physical anoxia that cells in the least mechanically reinforced milieus are most prone to anoxia-induced perturbations. Statistical threshold for identifying differentially-expressed genes across all pairwise comparisons was set as |log2FC| >= 1, i.e., 2-fold change in expression levels, with adjusted p-value < 0.05. (**h-m**) Ontological associations with biological processes for differentially-expressed genes, across oxo-mechanical combinations involving different degrees of chemical hypoxia induction and varying external mechanical milieus. (**n**) Heatmap comparing z-scores derived from relative gene expression levels for common targets of HIF1α, across varying external mechanical milieus (indicated by “0.3” and “1.5”, respectively, for 0.3 mg/ml and 1.5 mg/ml collagen matrices) for normoxia (indicated by “N”), anoxia (indicated by “A”), as well as chemically-induced hypoxia (indicated by the cobalt chloride dosage - 100 μM and 500 μM). (**o-q**) Representative maximum intensity projections of micrographs from cells embedded within 0.3 mg/ml, 1.5 mg/ml, and 2.5 mg/ml collagen matrices, exhibiting differential morphological responses to five different degrees of chemically-induced hypoxia. (**r-v**) Feret diameter measurements for cells embedded within five different collagen fiber density matrices (0.3, 0.5, 1.0, 1.5, and 2.5 mg/ml) and exposed to five different dosages of cobalt chloride (50, 100, 250, 500, and 1000 μM), with untreated cells (“Normoxia”) as a comparative control, showing how the individual cells in a population respond to chemically-induced hypoxia across distinctly different mechanical milieus. n > 100 individual cells for each condition, data represented as mean +/− s.d. Statistical significance calculated using an unpaired t-test, with significance marked at a p-value < 0.0001. (**w**) Phase space of cellular morphology defined by the external mechanical milieu (represented by collagen fiber density) and the intracellular oxygen state (represented by chemically-induced hypoxic signaling). All samples are compared between untreated and cobalt chloride-treated samples for each mechanical environment, with both being maintained under normoxia. Green squares represent similar morphologies between untreated and treated samples; red squares represent a more rounded-up population morphology within the treated samples as opposed to the more elongated population morphology within the untreated samples (p-value < 0.0001). These results demarcate a boundary for the oxo-mechanical effect to manifest, wherein, the increasingly reinforced external mechanical environment can proportionately buffer the morphology-perturbing effects of hypoxic signaling up until a threshold, beyond which the hypoxic stress dominates. Importantly, this establishes that the intracellular oxygen state directly alters the cell’s engagement with different mechanical milieus, thereby validating the concept of a reciprocal oxo-mechanical regulatory coupling.

Together, a distinct pattern of oxo-mechanics-dependent morphological states emerges. Specifically, we find that the least mechanically reinforced environment (0.3 mg/ml collagen matrix) and lowest oxygen availability (anoxia) together give rise to a morphologically-distinct population characterized by rounded-up cell bodies and low spread area (**Fig. 1t-1u**). However, normoxic conditions within the same mechanical regime enable cells to achieve elongated, spindle-like morphologies. By contrast, across every other tested oxo-mechanical regime, both anoxia and normoxia-exposed populations behave similarly, exhibiting elongated, spindle-like morphologies. These alterations in morphological behavior also manifest across prolonged incubation times and higher cell seeding densities (**SI Fig. 1c-1d**). Intriguingly, the effect of anoxia is strongly dependent on the mechanical microenvironment, as even the cells embedded within an incrementally higher fibre density matrix (0.5 mg/ml collagen) appear agnostic to reduced oxygen availability. To test this hypothesis across a broad range of oxo-mechanical regimes, we next expand our analyses over a combination of four different oxygen partial pressures and four distinct mechanical regimes (**Fig. 1v-1w** and **SI Fig. 4b-4f**). Herein, we only observe perturbed cellular morphology when both oxygen and mechanical reinforcement become limiting. Together, our combined morphometric analyses enable the first-such elucidation of an oxo-mechanical phase space capturing population-level cellular responses to simultaneously varying oxygen availability and environmental mechanics (**Fig. 1x**). Here, red squares represent conditions wherein both anoxic and normoxic populations maintained within the same mechanical regime differ in their behavior, whereas, green squares represent conditions of similar phenotypical manifestations. These data reinforce our findings that insufficient reinforcement from the external mechanical milieu allows low oxygen stress to profoundly affect cellular morphologies, whereas, reinforced mechanical regimes buffer the presumably deleterious effects of oxygen deprivation, suggesting a combinatorial oxo-mechanical regulation of cellular behavior.

### 2.2. A reciprocal oxo-mechanical coupling determines cellular states

To better describe the molecular signatures of oxo-mechanical regulation as well as characterize the cell states across different regimes, we perform bulk RNA-Seq using samples incubated under either anoxia or normoxia from both the least (0.3 mg/ml) and most (1.5 mg/ml) mechanically reinforced 3D collagen matrices. Across these, we visualize the differential gene expression profiles either as a consequence of altered oxygen availability (**Fig. 2a**), altered environmental mechanics (**Fig. 2b**), or a combination of both these variables (**Fig. 2c**). Interestingly, we capture the greatest extent of differential expression when both oxygen and mechanics are simultaneously altered, which is indicative of a combinatorial regulatory role (**Fig. 2a-2c**). More specifically, we also test for candidate groups of genes representing cellular adhesion, cytoskeletal components, ECM production, as well as hypoxic responses (**SI Fig. 5**). Given the widely-differing nature of these biological processes, a substantial degree of heterogeneity is expected across these samples, since changes to both the oxygen levels and environmental mechanics likely trigger a multitude of signalling pathways. Nevertheless, there is considerable value in exploring these transcriptomic datasets as a resource for understanding how combinatorial environmental cues alter both the overall cellular states as well as specific biological functionalities. To this end, we further perform ontology-based categorizations of the differentially-expressed genes and identify distinct classes of biological processes – such as adhesion, cytoskeletal function, cell cycle regulation, metabolism, and ECM production, amongst others – as being markedly altered upon variations in oxygen availability, environmental mechanics, as well as a combination of both (**Fig. 2d-2i**).

**Figure 5.**
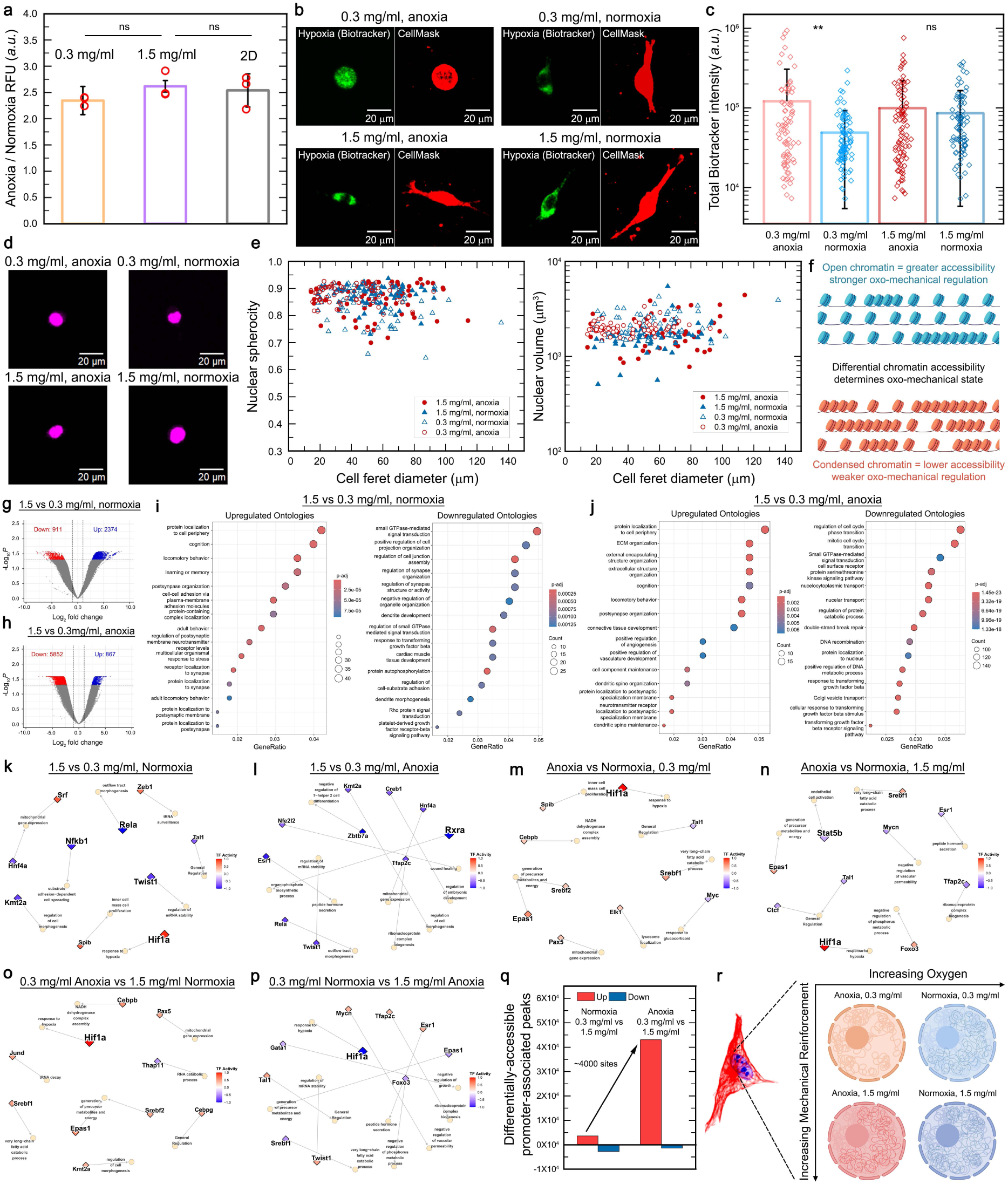
Differential chromatin accessibility controls oxo-mechanical regulation of cellular state. (**a**) Measurements of intracellular oxygen levels reveal no discernible changes due to either the collagen fiber density (0.3 mg/ml vs 1.5 mg/ml collagen) or nature of mechanical substrate (3D collagen matrices vs 2D plate). Hence, cells in both poorly-reinforced and well-reinforced collagen matrices experience similar levels of intracellular molecular oxygen for a given oxygen partial pressure across both mechanical regimes – hence, the input signal in terms of oxygen deprivation is agnostic to matrix mechanics. n = 3 independent replicates, data represented as mean +/− s.d. Statistical significance calculated using an unpaired t-test, with “ns” indicating p-value > 0.05. (**b** and **c**) Assessment of the intracellular hypoxic state using a reduction-activated fluorescent dye shows that cells embedded in poorly-reinforced mechanical regimes exhibit a significantly more hypoxic state under anoxia, in contrast to cells embedded in well-reinforced mechanical regimes, where differences between anoxia and normoxia-exposed cells are minimal. This suggests that the intracellular hypoxic state due to anoxic conditions is directly dependent on the external mechanical milieu. n > 75 individual cells for each condition, data represented as mean +/− s.d. Statistical significance calculated using an unpaired t-test, where “ns” indicates p-value > 0.05 and “**” represents p-value < 0.01. (**d** and **e**) Quantitative analyses of 3D nuclear morphology show no aberrant alteration of the nuclear shape due to specific oxo-mechanical regimes nor a dependence of nuclear morphology on the individual cell shape, exonerating perturbed nuclear architecture as the causative effect leading to rounded-up cell morphologies and altered cellular states. n > 55 individual cells for each condition. (**f**) Schematic representation of relaxed and compacted chromatin states – representing a potential mechanism for modulating regulatory control across different oxo-mechanical regimes. (**g-h**) Volcano plots for differentially-accessible chromatin loci, and (**i** and **j**) ontological association with biological processes for differentially-accessible promoter-associated chromatin loci, as determined by ATAC-Seq, compared between mechanically well-reinforced (1.5 mg/ml) and poorly-reinforced (0.3 mg/ml) external milieus across normoxia and anoxia. Statistical threshold for identifying differentially-accessible loci across all pairwise comparisons was set as |log2FC| >= 1, with adjusted p-value < 0.05. (**k-p**) Gene regulatory networks (GRNs) capturing the differential regulatory influence of transcriptional factors (TFs) across oxo-mechanical regimes. The diamond symbols represent specific TFs, whereas the connected circles represent the highest-ranking ontologically-classified biological process being regulated by its respective TF. The diamond size indicates the magnitude of TF regulatory influence, whereas, the diamond color indicates up/down-regulation with respect to the pairwise comparison. (**q**) ATAC-Seq analyses of promoter-associated genomic regions reveal that a combination of anoxia and poorly reinforced environmental mechanics retains the chromatin in a significantly more open and transcriptionally active state. (**r**) Together, these findings establish differential chromatin accessibility as the mechanism enabling microenvironments-specific oxo-mechanical regulation of cellular states.

Interestingly, we note that apart from a few overlapping functionalities, the biological processes being differentially expressed due to changes in both oxygen and mechanics (**Fig. 2h-2i**) are quite distinct from those which are triggered by changes to either one of these environmental variables (**Fig. 2d-2g**). This points towards a combinatorial regulatory framework, wherein, the simultaneous alteration to both oxygen and mechanics induce a unique set of alterations to the cell state, that cannot be otherwise captured by perturbing either of these variables in isolation. However, insights from RNA-Seq alone are a limited descriptor of cellular states, since we do not capture the contributions from post-translational regulation. Hence, to more comprehensively describe whether and how oxo-mechanical regimes alter cellular states, we next perform quantitative proteomics analyses on samples incubated under either anoxia or normoxia from both the least (0.3 mg/ml) and most (1.5 mg/ml) mechanically reinforced 3D collagen matrices. Here, we capture differentially-abundant proteins as a function of either altered oxygen availability, altered environmental mechanics, or a combination of both these variables (**SI Fig. 6**). Using these, we perform reactome analyses that identify specific biological pathways which are selectively enriched across different oxo-mechanical conditions – indicative of how these environmental regimes alter cellular functioning (**Fig. 2j-2o**). We further supplement this by performing ontology-based categorizations for biological processes (**Fig. 2p-2u**). Together, these analyses capture how biological processes such as adhesion, hypoxic responses, metabolism, cell cycle regulation, and ECM production are strongly influenced by a combination of environmental oxo-mechanical regimes (**SI Fig. 7**). Importantly, these findings also validate our results from the transcriptomic analyses – that a simultaneous variation in both oxygen and mechanics (**Fig. 2n-2o** and **Fig. 2r-2s**) manifests distinctly different biological effects than independent variations to either of these variables (**Fig. 2j-2m**, **Fig. 2p-2q**, and **Fig. 2t-2u**). Together, our interrogations at both the gene expression and protein expression levels suggests that varying oxygen partial pressures and altered external mechanical milieus profoundly influence cellular states in a combinatorial manner.

What do these altered cellular states imply for cellular functionality? From the reactome pathway analyses (**Fig. 2j-2o**), we observe distinct signatures corresponding to different oxo-mechanical regimes. Low oxygen availability induces metabolic reprogramming characterized by a glycolytic shift, as well as a lipogenic state, in both the poorly-reinforced (0.3 mg/ml anoxia) and well-reinforced (1.5 mg/ml anoxia) mechanical regimes (**Fig. 2j, 2k**, and **2n**). By contrast, cells in the poorly-reinforced matrix under normoxia (0.3 mg/ml normoxia) exhibit upregulation of oxidative phosphorylation (**Fig. 2l**) – which however, is not majorly upregulated in the well-reinforced matrix even under normoxia (1.5 mg/ml normoxia). This could potentially be linked to the pseudo-activation of HIF1a-associated responses (that promote a glycolytic shift) by increasing mechanical reinforcement even under normoxia, similar to previously reported cases^63–66^. Hence, the metabolic state of the cell is determined as a function of its local oxo-mechanical microenvironment. Interestingly, cells embedded in the well-reinforced mechanical milieus (1.5 mg/ml) in both anoxia and normoxia, exhibit an upregulation of integrin-based interactions (**Fig. 2l, 2n, 2m** and **2o**) – which helps explain why the morphological effects of hypoxic stress are prominently manifest only in the poorly-reinforced mechanical milieu (0.3 mg/ml anoxia). Finally, we also note that the cellular plasticity appears to be differentially potentiated by specific oxo-mechanical regimes. Specifically, we note that cells embedded in the poorly-reinforced mechanical regimes exhibit either a more transcriptionally active state (0.3 mg/ml normoxia) (**Fig. 2o**) or a high degree of chromatin remodelling (0.3 mg/ml anoxia) (**Fig. 2m**) – which are suggestive of a more plastic cellular state. Hence, these results collectively highlight that key aspects of the cellular morphology, metabolism, and regulatory program are differentially altered by environmental oxo-mechanical regimes – implying that the combination of oxygen partial pressures and external mechanical milieus determine cellular states.

Building on our findings from both the morphological characterizations and the altered cellular states (with regard to both gene and protein expression data), we now hypothesize that oxo-mechanical regulation manifests as a broad-spectrum reciprocal regulatory coupling. We posit that two broad tranches of regulatory components – the “oxygen state”, comprising all pathways related to sensing, perception, and response to oxygen levels; and the “mechanical state”, comprising all pathways related to adhesion, cytoskeletal activity, and mechanotransduction – interact as a reciprocal coupling (**Fig. 2v**). Specifically, this model implies that cellular states are an emergent property governed by the dialogue between two universal biological regulators – oxygen availability and environmental mechanics. To experimentally test this interpretation, we propose two distinct approaches – independently perturbing either the mechanical state or the hypoxic state and assessing the consequent response to either different oxygen partial pressures or different mechanical regimes, respectively.

### 2.3. Intracellular mechanical states determine cellular response to oxygen availability

To test the hypothesis that perturbing the cellular mechanical state alters response to oxygen, we systematically perturb two integral cytoskeletal components – the actomyosin contractility (using blebbistatin) and the microtubular stability (using colchicine). Here, we employ the highest non-lethal concentrations of blebbistatin (10 μM) and colchicine (500 pM) that ensure cell adhesion and elongation under normoxic conditions in the least mechanically reinforced regime (0.3 mg/ml – the softest and sparsest collagen matrix) (**SI Fig. 8**). A key experimental detail to note is that all comparative analyses are performed under identically-perturbed conditions across all oxo-mechanical regimes. Hence, any observed differences should be the consequence of oxo-mechanical regulation operating on cells with perturbed mechanical states. First, using blebbistatin treatments, we inhibit the actomyosin contractility, without significantly perturbing the actin organization in 3D, which shows a largely cortical localization that remains conserved irrespective of blebbistatin exposure (**Fig. 3a** and **SI Fig. 8a-8c**). Whereas previously, the effects of anoxia on altering the cell morphologies were only apparent in the least-reinforced mechanical milieu (0.3 mg/ml collagen), with the incorporation of blebbistatin treatment, this effect now prevails in the next highest mechanically reinforced regime (0.5 mg/ml collagen) (**Fig. 3b-3c**). By contrast, cells embedded in both the 1.0 mg/ml and 1.5 mg/ml collagen matrices exhibit elongated and spread-out morphologies under both anoxia and normoxia (**Fig. 3d-3e**), indicating that the combined perturbations exerted by blebbistatin treatment and anoxia can be countered only by significantly reinforced matrix mechanics (**Fig. 3f** and **SI Fig. 9a**). Effectively, cells with diminished actomyosin contractility require a higher degree of external mechanical reinforcement to overcome the effects of low oxygen availability.

Similarly, to perturb the microtubular stability, we employ a picomolar-range colchicine treatment, which appears to partially dissolve microtubules in 3D-adherent cells (**Fig. 3g** and **SI Fig. 8d-8f**). The treatment range is particularly critical, as even minor increments in colchicine-induced microtubular disruption significantly impedes adhesion of cells to 3D collagen matrices. Given this constraint, we systematically screen for the minimal colchicine dosage (1 nM colchicine) necessary to abolish adhesion in the least mechanically reinforced regime (0.3 mg/ml), and for all further experiments, maintain a treatment concentration just below this range (500 pM colchicine). In contrast to our experiments with blebbistatin treatments, we find that colchicine treatments when coupled with anoxia dramatically hamper cellular adhesion – rendering even cells in the most mechanically reinforced (1.5 mg/ml collagen) matrix with a rounded-up morphology (**Fig. 3h-3k**). Interestingly, this marked disruption does not occur as a result of the microtubular disruption alone. Rather, the coupled effects of destabilized microtubules and low oxygen availability together manifest a distinct effect on cellular morphology, giving rise to predominantly rounded-up cells (**Fig. 3l** and **SI Fig. 9b**). Compared against the data from cells without mechanical perturbation, the cells with targeted cytoskeletal perturbations exhibit a shift in the phase boundary – which delineates regimes exhibiting an oxo-mechanical effect on cellular morphology from regimes agnostic to low oxygen availability by virtue of reinforced environmental mechanics (**Fig. 3m**). Together, these findings validate that the intracellular mechanical state – exemplified by the cytoskeletal state here – profoundly alters the cellular response to different oxygen partial pressures.

### 2.4. Intracellular hypoxic signalling determines cellular engagement with external mechanical milieus

To test the hypothesis that perturbing the internal oxygen state alters cellular engagement with external mechanical milieus, we next perturb the intracellular hypoxic signalling. To directly modulate the intracellular hypoxic signalling, we employ a chemical mimetic of hypoxic signalling using the salt cobalt chloride (CoCl_2_)^67–70^, which stabilizes the hypoxia-inducible factor 1α (HIF1α), even under normoxic conditions (**Fig. 4a** and **SI Fig. 10a**). Further, controlling hypoxic induction through CoCl_2_ presents two distinct advantages. First, the effects of chemical hypoxia are dosage-dependent, which enables a graded administration of hypoxic treatments as verified by the transcriptomic analyses (**Fig. 4b-4c** and **Fig. 4e-4f**). Second, physical anoxia elicits readily discernible effects on cellular states only over a narrow range. For instance, morphometrically-discernible effects of anoxia – as captured by the oxo-mechanical phase space – appear constrained to the least-reinforced (0.3 mg/ml) 3D collagen matrix. Also, while anoxic conditions exhibit environmental mechanics-dependent differences in morphology compared to normoxia, these effects are ameliorated in regimes with higher mechanical reinforcement (>= 0.5 mg/ml). Using CoCl_2_ to induce hypoxic signalling overcomes this limitation, allowing us to investigate oxo-mechanical regulation over a much broader range of experimental regimes.

Similar to physical anoxia, both low (100 μM) and high (500 μM) dosages of CoCl_2_ elicit altered gene expression profiles which exhibit a marked dependence on the environmental mechanics (**Fig. 4b-4c**, **Fig. 4h-4i**, **Fig. 4e-4f**, **Fig. 4k-4l**, and **SI Fig. 10b-10e**). Precisely, for a given environmental mechanical regime, a strong induction of hypoxic signalling (high dosage of CoCl_2_) alters the expression of a greater number of genes when compared against normoxic samples (**Fig. 4e-4f**, **Fig. 4k-4l**), as opposed to a weak induction of hypoxia (low dosage of CoCl_2_) (**Fig. 4b-4c**, **Fig. 4h-4i**). The magnitude of this differential expression also depends on the external mechanical milieu, with larger perturbations being observed for cells within the poorly-reinforced (0.3 mg/ml) collagen matrix as opposed to the well-reinforced (1.5 mg/ml) collagen matrix (**Fig. 4b** vs **Fig. 4e**, **Fig. 4c** vs **Fig. 4f**). By comparison, keeping the degree of chemical hypoxia induction constant while varying the external mechanical regime does not dramatically alter the gene expression profiles (**Fig. 4d**, **4g**, **4j**, and **4m**). Hence, we conclude that cells embedded within the least mechanically reinforced matrix and subjected to strongly-induced hypoxic signalling exhibit the largest extent of differential gene expression, compared to their untreated normoxic counterparts (**Fig. 4e** and **Fig. 4k**). Conversely, cells embedded within the most mechanically reinforced matrix exhibit comparatively greater resistance to such perturbations (**Fig. 4f** and **Fig. 4l**). Together, these results establish that different degrees of oxygen state perturbation lead to variations in the cellular state depending on the external mechanical milieus. Furthermore, a direct comparison between transcriptomic signatures of HIF1α activity associated with either physical anoxia (environment oxygen partial pressure ∼ 0-0.2%) or CoCl_2_ treatments reveal very similar effects (**Fig. 4n**). Additionally, CoCl_2_ treatments also result in ontological terms associated with hypoxic signalling being identified as significantly upregulated – suggesting that CoCl_2_ indeed acts as a chemical inducer of hypoxic signalling (**Fig. 4h-4i**, **Fig. 4k**, and **Fig. 4n**).

Next, using single-cell morphometrics, we systematically assess how varying degrees of perturbed oxygen state alter cellular engagement with different mechanical environments (**Fig. 4o-4v** and **SI Fig. 11**). Combining five different dosages of cobalt chloride – 50, 100, 250, 500, and 1000 μM – with five different external mechanical milieus – 0.3, 0.5, 1.0, 1.5, and 2.5 mg/ml collagen fibre densities – we now quantify the cellular morphologies resulting from each such combination in comparison to the untreated normoxic control for each mechanical environment. While cells within the least mechanically reinforced collagen matrix (0.3 mg/ml) exhibit rounded-up morphologies following weak perturbations of oxygen state (100 μM CoCl_2_) (**Fig. 4o** and **Fig. 4r**), elevated mechanical reinforcement mitigates the effect of oxygen state perturbation up until a high dosage of CoCl_2_ (500 μM), at which stage cells invariably exhibit a rounded-up morphology, irrespective of the mechanical environment (**Fig. 4p-q** and **Fig. 4s-4v**). Pooling together data across these twenty-five distinct regimes, we describe a phase space wherein increasingly perturbed oxygen states directly counteract the stabilizing effects of increasing mechanical reinforcement (**Fig. 4w**). The phase space also highlights interesting parallels between the transcriptomic data and morphometric analyses. At low (100 μM) CoCl_2_ dosage, cells in the poorly-reinforced mechanical milieu significantly differ from those in the well-reinforced mechanical milieu, whereas, at high (500 μM) CoCl_2_ dosage, both poorly and well-reinforced mechanical milieus give rise to similar cellular states. These effects are in excellent agreement with trends observed in the transcriptomic data, wherein, the effect of perturbed intracellular oxygen signalling manifests in an environmental mechanics-dependent manner. Together, these data strongly validate that intracellular oxygen state – exemplified by the chemically-induced hypoxia here – profoundly alters the cellular engagement with external mechanical milieus. Furthermore, the oxo-mechanical phase space experimentally captures three distinct regimes of cellular responses – zones wherein the hypoxic signalling dominates, zones wherein the mechanical environment dominates, and intermediate zones wherein the competition between both determine cellular behavior. Hence, our findings support the model of a reciprocal oxo-mechanical coupling.

### 2.5. Differential chromatin accessibility controls oxo-mechanical regulation of cellular states

The coupling between a cell’s oxygen state and its mechanical state suggests that while response to varying oxygen levels is dictated by the underlying mechanical state, engagement with different mechanical regimes depends on the internal hypoxic signalling. Despite experimental data supporting this inference, a critical question remains unanswered – what is the mechanistic basis underlying this coupling? An immediate suspect is the cytoplasmic oxygen concentration – if the availability of molecular oxygen at the single cell level is set by the environmental mechanics, this would constitute a trivial explanation to the above question. To evaluate this, we employ a dye-based probe with oxygen-quenchable fluorescence – i.e., under high oxygen partial pressures, no signal is detected; whereas, low oxygen partial pressures give rise to fluorescent signal. However, our experiments comparing cells embedded within poorly-reinforced (0.3 mg/ml) and well-reinforced (1.5 mg/ml) 3D collagen matrices, as well as on 2D-adherent glass surfaces, across anoxia and normoxia do not capture any significant mechanical context-dependent differences in cytoplasmic oxygen levels (**Fig. 5a** and **SI Fig. 12a**). Rather, cells placed under anoxia appear to have similarly low internal oxygen partial pressures across all tested mechanical regimes in both 2D and 3D. Hence, oxo-mechanical phenomena are not a consequence of altered oxygen availability across different mechanical microenvironments – rather, these data suggest that different mechanical regimes likely alter how cells process the available oxygen. To test this, we use a cellular hypoxic state-reporting dye which undergoes a reduction reaction in the cytoplasm to generate a fluorescent product – hence, a more oxidative environment produces lower signal intensity, whereas a more reducing (hypoxic) environment gives rise to high signal intensity (**Fig. 5b**). Here, we quantify signal intensities from cells maintained under either anoxia or normoxia across both poorly-reinforced (0.3 mg/ml) and well-reinforced (1.5 mg/ml) collagen matrices. Remarkably, we find clear evidence of environmental mechanics-dependent hypoxic states. While cells in the poorly-reinforced mechanical milieu exhibit a significantly more hypoxic state under anoxic conditions, cells in the mechanically well-reinforced milieu do not exhibit any significant differences in their hypoxic state across both anoxia and normoxia (**Fig. 5c**). Hence, we conclude that poorly-reinforced environmental mechanics exacerbates the cell’s hypoxic state as opposed to well-reinforced environmental mechanics, which instead appear to buffer the effects of low oxygen availability. Together, these data suggest that a cell’s oxygen processing is directly linked to its external mechanical milieu.

These observations raise the question – what mechanisms do cells employ to perceive their external oxo-mechanical regimes and accordingly modulate internal cellular states? Given the wide range of oxo-mechanical regulatory effects evident from both morphometric analyses and transcriptomic/proteomic profiles – none of which implicate a single signalling pathway operating in isolation – it is conceivable that a broadly influential physical mechanism underlies these phenomena. Motivated by quantitative measurements of both cellular oxygen levels and hypoxic states pointing towards a mechanics-driven influence over oxygen processing, we now consider one such broad-acting effect of substrate mechanics – altered chromatin states. Prior work provides compelling evidence demonstrating how altered substrate mechanics significantly alters chromatin accessibility^71–74^. Furthermore, from our proteomics analyses (**Fig. 2o** and **2m**) we note that cells embedded in poorly reinforced mechanical milieus (0.3 mg/ml) exhibit a more plastic state – as evidenced by either higher transcriptional activity (0.3 mg/ml normoxia) or increased chromatin remodelling activity (0.3 mg/ml anoxia). Given that oxo-mechanical effects are most profoundly manifested by cells embedded within poorly-reinforced low fibre density collagen matrices under anoxia, as opposed to cells embedded within well-reinforced high fibre density collagen matrices under anoxia, we hypothesize that similar mechanisms could be at play here. Precisely, altered genomic accessibility mediated by environmental mechanics could enable similar levels of hypoxic signalling-induced transcriptional factors to elicit dramatically different phenotypical effects.

To investigate this possibility, we first compare the nuclear morphologies of cells maintained under either anoxia and normoxia across both poorly-reinforced and well-reinforced collagen matrices (**Fig. 5d**). Interestingly, nuclear shapes do not exhibit any specific trend suggestive of oxo-mechanical regulation (**Fig. 5e** and **SI Fig. 12b-12c**). This is especially striking, considering that the overall cellular morphology does recapitulate the prior observations of being predominantly rounded-up under low oxygen and poorly reinforced mechanical regimes. Indeed, we find no clear correlation between the overall cell morphology and the nuclear morphology – suggesting that in these 3D-adherent cells, these two aspects are likely decoupled from each other. However, timelapse imaging suggests that the nuclei in these 3D-adherent cells are not static entities. Rather, these appear to undergo discernible shifts in their overall shapes as well as in the distribution and spatial localization of sub-nuclear speckles, which may likely alter the chromatin arrangements within the nuclei, potentially leading to altered gene expression profiles (**SI Fig. 12d** and **SI Video 5**). Importantly, despite a lack of oxo-mechanical influence over ensemble nuclear shapes, such intra-nuclear plasticity could allow altered environmental mechanics to modify chromatin conformations (**Fig. 5f**).

We test this hypothesis by globally characterizing the chromatin states in cells maintained under either anoxia or normoxia within both poorly-reinforced and well-reinforced collagen matrices. Using ATAC-Seq, our analyses capture population-level profiles for genomic loci retained in open conformations across these oxo-mechanical regimes. Interestingly, we find that a change in oxygen partial pressure alone (normoxia vs anoxia) does not lead to any significantly differentially-accessible chromatin loci, regardless of the external mechanical milieu. However, altering the external mechanical milieu dramatically changes the differential accessibility of several thousand chromatin sites in both normoxic and anoxic conditions – with key differences between these two combinations. Under normoxic conditions, a larger number of sites appear accessible in the well-reinforced mechanical milieu as opposed to the poorly-reinforced mechanical milieu (**Fig. 5g**). However, under anoxic conditions, this pattern reverses, as, a much larger number of sites becomes accessible in the poorly-reinforced mechanical milieu as opposed to the well-reinforced mechanical milieu (**Fig. 5h**). Interestingly, ontology-based annotation of the identified chromatin loci with biological processes reveals that the differentially-accessible loci in both comparisons (**Fig. 5i-5j**) are also associated with several functionalities that remained previously unidentified in our transcriptomic analyses. Together, these results suggest that differential chromatin accessibility may be leveraged to orchestrate oxo-mechanical regulatory programs under specific combinations of environmental regimes.

To develop a more comprehensive description of the regulatory framework controlling cellular state across oxo-mechanical regimes, we now integrate both the differential gene expression data from RNA-Seq as well as the differential chromatin accessibility data from the ATAC-Seq to define gene regulatory networks (GRNs). Using a GENIE3 machine learning-based approach, we first identify transcription factors (TFs)-gene target interactions from the RNA-Seq data. We evaluate the regulatory influence of these TFs (**SI Fig. 13**) by incorporating TF binding evidence using motif matching within accessible chromatin peaks, estimates of TF activity, as well as TF-target correlation. We identify the top ten such TFs exhibiting the largest regulatory influence for each pairwise comparison, as well as their top hundred targets, which are subsequently subject to an ontology-based classification to identify the specific biological process being differentially regulated. We have additionally included a complete list detailing the TF regulatory scores and target scores (**SI Table 1 – SI Table 13**), as well as an overall GRN spanning all four oxo-mechanical regimes (**SI Fig. 14** and **SI Table 14**). For clarity, however, we restrict our graphical representation of GRNs describing the regulatome of each pairwise comparison across variations in oxygen partial pressures, mechanical reinforcement, or both variables together, to the highest-scoring biological process for each individual TF (**Fig. 5k-5p**).

Under anoxic conditions, irrespective of external mechanical milieus, we observe strong signatures of HIF1α and Epas1 (HIF1β)-driven regulation. This likely manifests in the form of metabolic reprogramming via glycolytic shifts, lipogenesis, and cholesterol biosynthesis. This is most prominently observed in poorly-reinforced mechanical environments under anoxia (0.3 mg/ml anoxia) by the upregulation of both Srebf1 and Srebf2 (**Fig. 5m**), as well as to a lesser extent in well-reinforced mechanical environments under anoxia (1.5 mg/ml anoxia), wherein Srebf1 is upregulated. Interestingly, anoxic conditions manifest very different cell states between poorly-reinforced and well-reinforced external mechanical milieus – wherein, enhanced mechanical reinforcement significantly reduces the TFs activity under anoxia. Specifically, each of the top ten TFs in the comparison between 1.5 mg/ml anoxia vs 0.3 mg/ml anoxia (**Fig. 5l**) exhibit downregulation in the former (thereby upregulation in the latter). This implies that well-reinforced mechanical regimes (1.5 mg/ml) make the cell state less plastic (more inaccessible / lower TFs regulatory influence), whereas, poorly-reinforced mechanical regimes (0.3 mg/ml) retain the cells in a much more plastic state (more accessible / higher TFs regulatory influence). This difference has direct consequences for cellular adaptation to hypoxic stress, whereby, less-reinforced mechanical milieus enable a broader range of stress-adaptive strategies. We observe that the primary hypoxic stress adaptation signature in well-reinforced matrices (1.5 mg/ml anoxia, **Fig. 5n**) is an upregulation of Foxo3 – a broadly-acting stress response pathway, which is also HIF1α-inducible. By contrast, cells in poorly-reinforced mechanical milieus (0.3 mg/ml anoxia, **Fig. 5l** and **5o**) utilize multiple such adaptive responses – Jund (cell survival), Nfe2l2 (oxidative stress regulator), Creb1 (cell survival), Rxra (metabolic reprogramming via enhanced glycolysis), and Twist1 (apoptosis suppression). Hence, a more accessible chromatin state in poorly-reinforced milieus (0.3 mg/ml) enables a larger regulatory influence manifested by multiple stress adaptive programs against anoxia, whereas, less accessible chromatin in well-reinforced milieus (1.5 mg/ml) limits the cell’s options in response to anoxia. Furthermore, we also note that cellular proliferation is significantly reduced under anoxic conditions across both poorly and well-reinforced mechanical milieus – as shown by downregulated Myc/Mycn levels (**Fig. 5m** and **5n**) – whereas, under poorly-reinforced normoxic conditions cells appear to retain a proliferative state via upregulation of Mycn (**Fig. 5p**).

It is also interesting to note that oxo-mechanical regimes appear to enable parallel paths towards accomplishing similar phenotypic outcomes. For instance, cells embedded in both poorly-reinforced (0.3 mg/ml) and well-reinforced (1.5 mg/ml) mechanical regimes under normoxia demonstrate an upregulation of mesenchymal phenotypes (**Fig. 5k**). However, in the case of cells within poorly-reinforced milieus under normoxia (0.3 mg/ml normoxia), the mesenchymal state is likely driven by the Rela/Nfk-β signalling axis acting via Twist1 induction. By contrast, in the well-reinforced milieus under normoxia (1.5 mg/ml normoxia), the mesenchymal state likely comes about as a consequence of the HIF1α-targeted Zeb1 activity. It is worth noting that both Twist1 and Zeb1 are also central players in the EMT transition, which promotes more mesenchymal and migratory phenotypes. Along similar lines, we also find that how strongly a particular cell state is induced depends on the particular oxo-mechanical cue responsible for triggering the underlying regulatory state. This is exemplified by the apparent pseudo-activation of HIF1α by an increase in mechanical reinforcement under normoxic conditions (**Fig. 5k**). In all other cases, the upregulation of HIF1α requires low oxygen partial pressures. The mechanically-driven activation of HIF1α has prior evidence in the literature and has also been shown to be associated with ECM remodelling functionalities driven by modulators such as Lox, Plod2, Mmp2, and P4h41/2^75,76^. This possibility is also supported by the proteomics analyses, wherein, cells embedded in the well-reinforced mechanical environment exhibit an upregulation of ECM remodelling functions (**Fig. 2l, 2m, 2n**, and **2o**). However, we also highlight that a comparison between well-reinforced and poorly-reinforced mechanical milieus under anoxia does not exhibit an upregulation of the HIF1α signalling (**Fig. 5l**). This discrepancy suggests that mechanically-driven activation of HIF1α occurs to a much lesser extent than low oxygen partial pressures-driven activation of HIF1α. Together, these results highlight that similar biological signalling pathways may be separately inducible by very different environmental cues.

The inferences from the GRN analyses also help explain our initial morphological observations. A comparison between cells embedded in well-reinforced mechanical milieus (1.5 mg/ml) and poorly-reinforced mechanical milieus (0.3 mg/ml) under normoxic conditions (**Fig. 5k**) reveals that cells in the former assume a more mechanically-activated and contractile state, as inferred by the upregulation of Srf, that is critical for the regulation of cytoskeletal genes (especially actin/actomyosin) and has positive associations with focal adhesion complexes necessary for adhesion. These results are also in agreement with our proteomic analyses suggesting that integrin-associated interactions are enriched for cells grown in the mechanically well-reinforced matrices (**Fig. 2l, 2m, 2n**, and **2o**). Together, these observations help explain the trends observed with our morphometric analyses – wherein, cells embedded within the poorly-reinforced matrix are more detrimentally affected by anoxic stress (manifested by low spread area and rounded morphologies) as opposed to cells embedded within the well-reinforced matrix. It is likely that the stress due to hypoxic responses is buffered by the upregulated Srf activity in the well-reinforced matrices, thereby, enabling the cells to retain a spread-out elongated morphology even under anoxia. However, we also note that a comparison between cells in the well-reinforced vs poorly-reinforced matrices under anoxia does not exhibit a differential regulation of Srf (**Fig. 5l**) – which could be a consequence of HIF1α activity counteracting the mechanical stabilization. Importantly, when the hypoxic stress due to anoxia is further exacerbated by either a cytoskeletal perturbation (colchicine treatment, **Fig. 3m**) or chemically-induced hypoxia (cobalt chloride treatment, **Fig. 4w**), we find that cells assume rounded-up morphologies even in the well-reinforced matrices – suggesting that while the stabilizing effects of Srf are countered by anoxia, any further destabilization of the cells due to additional cytoskeletal or hypoxic perturbations pushes them towards a rounded-up morphology. Hence, we conclude that the influence of external mechanical milieus alters both the contractility and mechanical activation of cells, whereas, the influence of low oxygen partial pressures further modulates the extent of intracellular mechanical stabilization.

In relation to chromatin accessibility, we also find further evidence supporting mechanically-tuned alteration of chromatin organization from the GRN analyses. In particular, we note that the chromatin modifier Kmt2a is upregulated in poorly-reinforced milieus when compared against well-reinforced milieus, across both anoxic and normoxic conditions (**Fig. 5k** and **5l**). However, a change in oxygen partial pressures does not significantly alter the differential activity of Kmt2a (**Fig. 5m** and **5n**). Interestingly, upon comparing cells from poorly-reinforced milieus under anoxia against cells from well-reinforced milieus under normoxia, we find a distinct upregulation of Kmt2a (**Fig. 5o**). However, and critically, no comparable differential activity is observed when comparing cells from poorly-reinforced milieus under normoxia against cells from well-reinforced milieus under anoxia (**Fig. 5p**). Together, these findings suggest that while an alteration in oxygen partial pressures does not appreciably alter the chromatin accessibility, and an alteration in the mechanical reinforcement significantly alters the chromatin accessibility, it is a coupled change in both oxygen partial pressures and the external mechanical reinforcement that together manifest a pronounced difference in the cell’s chromatin remodelling functions – as further supported by our direct quantification of promoter-associated accessible chromatin peaks from the ATAC-Seq below (**Fig. 5q**). Hence, we infer that cellular plasticity is differentially highest under the oxo-mechanical regime corresponding to poorly-reinforced anoxic regimes (0.3 mg/ml anoxia) – which thereby represents the most accessible context for oxo-mechanical regulatory effects to manifest.

Along with our transcriptomics and proteomics analyses, the above observations suggest a putative role for different oxo-mechanical regimes to potentiate functionally-distinct fibroblast forms. Using our GRNs analyses, we can now broadly categorize cells across the four oxo-mechanical regimes being interrogated. First, the poorly-reinforced mechanical milieus under anoxia induce highly plastic fibroblasts states, which are characterized by quiescence (lack of proliferation), low mechanical activation and contractility, distinct metabolic reprogramming via glycolytic shifts and lipogenesis, a robust adaptive response to hypoxic stress, as well as a high degree of TFs regulatory influence coupled with a strong upregulation of chromatin remodelling. Second, the poorly-reinforced mechanical milieus under normoxia induce inflammatory fibroblasts states, which are characterized by a high Rela/Nfk-β inflammatory activity, upregulation of proliferative signals, as well as poor mechanical activation and contractility. Third, the well-reinforced mechanical milieus under anoxia induce fibrotic fibroblast states, which are characterized by quiescence (lack of proliferation), high mechanical activation and contractility, partial metabolic reprogramming via glycolytic shifts and lipogenesis, as well as a broad adaptive response to hypoxic stress. Fourth, the well-reinforced mechanical milieus under normoxia induce proto-myofibroblast states, which are characterized by a high degree of mechanical activation and contractility, presence of proliferative signals, mechanical activation of HIF1α, as well as an elevated potential for ECM remodelling. Taken together, these results suggest that different oxo-mechanical regimes modulate cellular regulatory programs to varying extents, based on the specific combination of oxygen partial pressures and external mechanical milieus.

What is the mechanistic basis for these differentially-activated regulatory programs? How do specific combinations of oxygen partial pressures and mechanical milieus effect dramatic alterations to almost all aspects of cellular physiology – ranging across morphology, gene expression profiles, protein expression, and regulatory modules? Motivated by the observation that oxo-mechanical regimes differentially regulate chromatin accessibility, we next compare the proportion of differentially accessible chromatin loci annotated as promoter-associated regions (suggesting a likelihood for transcriptional regulation) across different environmental regimes. Here, we discover that transcriptional accessibility is directly controlled by environmental oxo-mechanical regimes (**Fig. 5q**). While variations in oxygen partial pressures alone do not appear to significantly affect chromatin states, varying the external mechanical milieu differentially alters the accessibility of promoter-associated chromatin loci. Even under normoxic conditions, there is only a modest difference between cells embedded in mechanically poorly-reinforced and well-reinforced collagen matrices. However, a combination of low oxygen and poor mechanical reinforcement remarkably enhances chromatin accessibility for promoter-associated regions – compellingly demonstrating a coupled oxo-mechanical regulation. Effectively, anoxia and poorly-reinforced mechanical milieus together render cells significantly more responsive to oxo-mechanical perturbations, which is also reflected in their morphological behavior and TF regulatory influence. Therefore, we propose that oxo-mechanical environments-determined differential chromatin accessibility enables and modulates a coupled regulation of cellular states (**Fig. 5r**). Importantly, we emphasize that these effects cannot be recapitulated by either of these environmental variables in isolation. Rather, only specific combinations between oxygen and mechanics lead to such emergent behaviors and cell states, underscoring the idea of a coupled oxo-mechanical regulation.

## 3. Discussion

Our work presents a coupled regulatory framework between oxygen and mechanics, which governs cellular states in 3D ECM-like microenvironments. Across diverse biological scales and species, oxygen availability and environmental mechanics represent two of the most influential physical agents, with broadly-acting effects on physiology, metabolism, and organization^4,13,14,18,44,53,55^. Whereas prior work has extensively characterized their effects as independent variables, we advance a unique perspective on how both oxygen and mechanics act in tandem to exert a combinatorial regulation of cellular states. Importantly, we establish that cellular perception and response to oxygen partial pressures is determined by intracellular mechanics, whereas, cellular engagement with varying external mechanical milieus is determined by intracellular hypoxic signalling. Further, we identify differential chromatin accessibility as the mechanistic basis for these phenomena – wherein, specific combinations of oxygen partial pressures and external mechanical milieus control the regulatory influence exerted by transcriptional factors towards determining distinct cellular states. Taken together, our work establishes oxo-mechanical regulation as a fundamental paradigm of cellular sensing and response in complex 3D microenvironments, as well as demonstrates how cellular states are an emergent property of their surroundings. Importantly, the definition of an oxo-mechanical framework paves the way towards exciting new questions and future investigations.

A key feature observed across our transcriptomic, proteomic, chromatin accessibility, and regulatome (GRN) analyses is the pronounced enrichment of elements associated with reprogramming and pluripotency. This directly suggests that oxo-mechanical regimes regulate cellular plasticity – which is intriguing considering that prior work shows how each of oxygen and substrate mechanics act as critical determinants of cell fate transitions. However, tissues – in particular, stem cell niches - are dynamic in nature, wherein both oxygen availability and environmental mechanics exhibit spatiotemporal variations^17,18,44,46,48–50^. Against this backdrop, it becomes interesting to ask how the local microenvironment around a stem cell population either potentiates or restricts prospective fate transitions. This has considerable value towards understanding the physical principles underlying stem cell niche design as well as processes ensuring the maintenance of stem cell pools in vivo while simultaneously allowing for controlled differentiation. Furthermore, our work demonstrates signatures of reprogramming-associated modules being differentially regulated in fibroblast cells purely as a function of their external environment. This is intriguing from a translational angle – specifically with regard to in vitro strategies that employ chemical cocktails to induce pluripotent states in differentiated cells^77–80^. It is conceivable that specific combinations of oxygen and mechanics modulate the checkpoints gating transitions between cell states, which could either enable or restrict fate transition and dedifferentiation processes. Hence, exploring the combinatorial role of both oxygen and mechanics in stem cell biology will be of both fundamental and translational interest.

The complementary perspective to how microenvironments regulate cell states - i.e., how cells remodel their local microenvironments – also brings up interesting considerations in the context of oxo-mechanical coupling. Fibroblasts are key players in cell-driven niche remodelling, and prior work has extensively characterized how mechanical perturbations (altered substrate stiffness / contractility) as well as oxygen stress triggers the emergence of myofibroblast phenotypes that remodel the ECM to maintain tissue homestasis – such as during wound repair, wherein, loss of regulatory control over this process leads to scarring^81–87^. Whereas the experiments in our study are almost exclusively restricted to short time scales (∼24 hours) and very low cell densities, it is likely that over extended durations and higher cell packing fractions approaching those of solid tissues, the induction of such phenotypes under different oxo-mechanical regimes may substantially alter tissue architecture and mechanical integrity. Conversely, how different combinations of oxygen partial pressures and external mechanical milieus alter the kinetics of phenotypic transitions also remain to be explored. As noted in our transcriptomic and proteomic analyses, ECM production, organization, and remodelling prominently feature amongst the class of biological processes that are highly responsive to oxo-mechanical regimes. This aspect is valuable towards understanding wound healing processes, wherein, the balance between controlled and excess ECM production crucially determines whether tissues remain intact or turn fibrotic. Wound healing presents an intriguing scenario with regard to oxo-mechanical regulation, given the confluence of diverse cell types (fibroblasts, neutrophils, and macrophages), local hypoxia (due to vascular damage and/or induction of inflammatory responses), and variable microenvironmental mechanics (loss of tissue integrity followed by clotting and induction of repair processes including ECM deposition)^88–95^. The spatiotemporally dynamic and highly interlinked nature of these processes presents both a complex modelling challenge as well as a rich arena for understanding how the interactions between cells and their local oxo-mechanical microenvironment determines tissue fate – especially using engineered in vitro 3D models, wherein both oxygen availability as well as global/local mechanics can be precisely manipulated.

Our understanding of oxo-mechanical regulation also necessitates a more nuanced definition of the term “hypoxia”. Prior work has demonstrated that hypoxic responses are very cell type-specific – indeed, deprivation of oxygen manifests both marginal and massively detrimental effects^18,44,46,96–99^. However, our findings suggest that cellular perception and response to oxygen availability is not a function of the oxygen partial pressure alone. Rather, the cell’s external mechanical milieu and its internal mechanical state are key determinants of how cells perceive and respond to reduced oxygen availability, as well as the extent to which hypoxic responses are activated. Conversely, the extent to which intracellular hypoxic signalling is activated directly controls the cell’s engagement with its external mechanical environment. This coupling advocates for a more context-sensitive definition of cellular hypoxia – which could also help explain prior observations of cell type-specific effects exerted by reduced oxygen partial pressures. Importantly, this perspective also suggests that tissue remodelling by ECM deposition in hypoxic niches is, at least in part, an adaptive response by cells to reinforce their environmental mechanics and thereby render themselves more robust against low oxygen stresses. This also adds an important dimension to our findings suggesting that cellular states are emergent properties of their local microenvironments, by now involving an active cell-driven component of microenvironmental remodelling in response to the original environmental regime.

Our results also raise an elementary, albeit challenging, question – how will the patterns of oxo-mechanical regulation manifest for different cell and tissue types? Our present work employs fibroblasts – differentiated mesenchymal cells – using which, we have defined phase spaces spanning different sets of perturbations, as well as molecular descriptions of cellular states across different oxo-mechanical regimes. However, it is likely that a biologically-distinct system – such as cancer cells, which feature severely dysregulated hypoxic signalling and intracellular mechanics^11,18,51,100,101^ – may dramatically alter the contours of these phase spaces as well as only exhibit discernible differences in regimes that lie outside the boundaries explored in this work. Furthermore, cancer cells exhibiting collective organization in form of spheroids and tumors intrinsically feature elements of oxo-mechanical heterogeneities due to their 3D spatial organization, which sets up a natural gradient of mechanical compression and oxygen limitation originating from the periphery and extending into the core^102–107^. Hence, within the same ensemble structure, different cellular states would be expected to emerge in a spatially-defined fashion – potentially contributing towards phenotypic single cell-level heterogeneity within a clonally-identical population. Finally, we also note that major classes of differentially-regulated processes identified by our multi-omics analyses – such as cell cycle regulation, metabolism, and reprogramming – typically manifest in a highly cell type-specific manner. Even though the fundamental workings of oxo-mechanical regulation employ generic mechanisms – ubiquitous environmental variables, conserved biochemical signalling modules, and large-scale chromatin remodelling – it is highly probable that the specific outcomes of these effects will be heavily influenced by the biological identity of the system being interrogated. Consequently, there is a need to engineer versatile 3D culture platforms that can effectively accommodate multiple such model systems. The design principles for these could incorporate – functionalizable 3D matrices featuring different classes of adhesion-enabling moieties^108^; non-adherent 3D milieus (especially for immune cells)^109^; mechanical tunability over a broad range of biomimetic regimes^110^; adjustable cell densities without compromising material properties; modular nutrient chemistry and growth factor availability^111,112^; as well as capabilities to accommodate both single cells and multicellular collectives^113–115^.

The oxo-mechanical coupling presented in this work is particularly potent because it integrates two of the most conserved, ubiquitous, and influential environmental forces that constantly affect almost all scales and forms of living systems. We intend this as a bedrock for guiding future investigations across cells, multicellular collectives, and model organisms, with the eventual goal of better understanding the myriad ways in which physical microenvironments actively regulate their resident living matter.

## 4. Materials and Methods

### 4.1. Preparation, rheological characterization, and imaging of 3D collagen matrices

We prepare 3D ECM-like matrices using collagen type I sourced from rat tail (Gibco), supplied as a 3 mg/ml stock solution. To prepare collagen matrices with different collagen fiber densities we utilize a reaction mixture comprising 10X DMEM, 0.1 N NaOH (used at ¼ volume of collagen stock solution), 3 mg/ml collagen type I monomer solution, and dilute cells’ suspension (final concentration <100 cells per μL for all experiments, unless specified otherwise for limited experiments), with distilled water used to make up the final volume. The amount of collagen monomer solution added is adjusted to achieve final concentrations of 0.3 mg/ml, 0.5 mg/ml, 1.0 mg/ml, 1.5 mg/ml, and 2.5 mg/ml. Immediately after preparation, the mixture is incubated at 37°C for a maximum of 45 minutes to allow complete polymerization. Following this, we add fresh cell culture media, and house the samples under the appropriate environmental oxygen concentration as per the required experiment. This strategy ensures that cells are homogeneously dispersed throughout the bulk matrix prior to polymerization itself. The collagen fiber density in such 3D matrices can be directly visualized using reflectance microscopy which captures the reflections of individual fibers as high-contrast elements – for these imaging experiments alone we prepare the matrices without adding cells. We also ascertain the viscoelastic properties of these collagen matrices using shear rheology with an Anton Parr 302e rheometer. Briefly, the reaction mixtures for different collagen density matrices are prepared sequentially and polymerized at 37°C for 45 minutes while sandwiched between the base plate and a roughened cone plate. During this time, we maintain high environmental humidity to prevent sample desiccation. Following this, we apply small amplitude (1%) shear while recording the complex shear moduli for 10 minutes. This value is further resolved into components of elastic storage modulus (G’) – indicating the energy stored in the material during a deformation cycle, and thereby, its elastic solid-like nature – and viscous loss modulus (G”) – indicating the energy dissipated by the material during a deformation cycle, and thereby, its viscous fluid-like nature. Since G’ > G” for all samples, we confirm that all collagen matrices are stable solid structures following 45 minutes of polymerization at 37°C.

### 4.2. Cell culture

For all experiments we employ the immortalized NIH-3T3 (mouse embryonic fibroblasts) cell line, with cells being used only between the third and tenth passages. Cell culture media comprises of DMEM supplemented with 10% FBS, 1% Pen-Strep, sodium pyruvate, and the sodium bicarbonate buffering system. All cell culture is performed with 5% environmental CO_2_ partial pressure, with the oxygen partial pressures being varied between ∼0.2% (anoxia) and ∼20% (normoxia), as well as intermediate values of ∼3% and 10% for a limited series of experiments. To alter oxygen levels, we use a tri-gas mixer (OkoLab) equipped with a system of three parallel flow meters to regulate compressed air, CO_2_, and N_2_ levels.

### 4.3. Imaging and analysis of 3D cell morphology

All images are acquired using a point-scanning laser confocal microscope (Nikon A1HD25 Ti2 Eclipse). All 3D collagen matrices for imaging-based experiments are prepared in glass-bottom 35 mm dishes. Prior to imaging, we stain cells with calcein-AM for atleast 30 minutes. For single-cell morphometrics, we acquire >100 cells per experimental sample type as individual z-stacks spanning the entire cell volume. All data is acquired at least 12 hours following sample preparation to allow for cells to adhere and spread in the 3D collagen matrices. We generate maximum intensity projections of all such z-stacks and manually outline the cell body for each such image using ImageJ, following which we use in-built particle analysis functions to extract quantitative measurements of morphological parameters. For principal component analysis we use custom MATLAB scripts which take the population average values of seven different morphological parameters for all eight sample types. Similarly, we use MATLAB to convert raw population-level data from each sample type into appropriate probability distributions using both the probability density function and complementary cumulative density function.

### 4.4. Imaging and analysis of nuclear morphology

To characterize the nuclear morphology of cells embedded in 3D collagen matrices, we prepare samples as described above, and prior to imaging, co-stain with calcein and Hoechst for 30 minutes. Following this, we acquire z-stacks of individual cells to capture both the cell body as well as the nucleus. Using custom MATLAB scripts, we first interpolate the z-stack data to reconstruct a 3D mesh that conserves the actual object dimensions, followed by 3D segmentation to demarcate the nuclear region alone and obtain morphometric parameters describing its conformation. We also perform similar morphological analysis using maximum intensity projections of the z-stack images to compare between both modes of analysis. Finally, we manually segment the cell body corresponding to each such nucleus in ImageJ and measure its feret diameter, which is then compared against the nuclear morphology.

### 4.5. Measurement of cell viability

To assess the effect of both physical anoxia as well as chemically-induced hypoxic signalling on cell viability, we use calcein-AM (to mark live cells) and propidium iodide (to mark membrane compromised, presumably dead, cells). Cells pre-stained with calcein-AM are maintained in the 3D collagen matrix either under normoxia or anoxia for 24 hours, following which we stain the cells with propidium iodide for 15 minutes. Images are acquired as a confocal z-stack, and maximum intensity projections of these are used to tally the number of live and dead cells. All cells which only show calcein-specific fluorescence (green) are assumed to be viable, whereas, those which show both calcein and propidium iodide-specific fluorescence (green + red = yellow) are assumed to be dead.

### 4.6. Drug treatments - blebbistatin, colchicine, and cobalt chloride

To determine the specific concentrations of blebbistatin and colchicine used for perturbing the intracellular mechanical state, we systematically screen through several treatment dosages starting with values typically reported in the literature and apply these to cells in 0.3 mg/ml collagen matrices incubated under normoxic conditions. The criteria here is to find the highest such concentration wherein cellular spreading and elongation under normoxic conditions in the least mechanically reinforced collagen matrix (0.3 mg/ml) is not drastically perturbed. We thus identify a concentration of 10 μM for blebbistatin and 500 pM for colchicine. We visualize the intracellular actin and microtubule organization using the live-staining dyes sir-Actin and sir-Tubulin, respectively (obtained from Cytoskeleton, Inc.). For cobalt chloride, we first identify that previous work reports concentrations in the range of 1-10 mM for exerting a chemical hypoxia signalling effect on cells. Hence, we perform assays across a range of concentrations ranging from 50 μM to 1 mM of cobalt chloride treatments provided to cells under normoxic conditions.

### 4.7. Measurement of intracellular oxygen levels and reducing/oxidative state

For measuring the intracellular oxygen concentration, we use a proprietary time-resolved fluorescence-based dye (Abcam, ab197245), which is quenched by presence of molecular oxygen. We pre-load cells with this dye and seed them on either collagen monomers-coated (a 10 μg/ml solution used to coat for 30 mins at room temperature) 2D glass surfaces, or within both 0.3 mg/ml and 1.5 mg/ml 3D collagen matrices. Using a multimode plate reader (VarioSkan Lux, ThermoFisher) operating in the time-resolved fluorescence mode, we conduct a spectrum scan across the entire recommended operating range for the dye and calculate the ratio between signal captured from anoxic and normoxic samples for all three conditions for the excitation-emission wavelength combinations showing optimal signal generation across all conditions. To evaluate the intracellular reducing/oxidative state, we use the BioTracker 520 Green Hypoxia Dye to stain cells embedded in 3D collagen matrices prepared as described above, 12 hours post-incubation under the appropriate oxygen partial pressures. We also co-stain the cells with CellMask Red to segment the cell boundary and acquire z-stacks of individual cells as described above. We trace the cell outlines in ImageJ and quantify the sum-slices signal intensity for the BioTracker 520 Green Hypoxia Dye to measure the intracellular reducing/oxidative state – a higher value indicates a more reducing intracellular environment, whereas a lower value indicates a more oxidative intracellular environment.

### 4.8. Immunostaining of HIF1α

Samples for immunostaining are prepared by seeding cells on glass-bottom dishes and incubating under either normoxia, anoxia, or normoxia + 100 μM CoCl_2_ / 500 μM CoCl_2_. We fix these samples using 4% PFA for 15 minutes at room temperature, followed by three washes in PBST buffer (0.2% Tween-20 in PBS). The fixation reaction is quenched using 0.15 M glycine buffer by incubating for 15 minutes at room temperature, followed by three washes in PBST buffer. The samples are permeabilized using a 1% Triton X-100 solution in PBS, for 1.5 hours at room temperature, and further blocked with 1% BSA for 6 hours at room temperature, followed by three washes in PBST buffer. We perform primary antibody (anti-HIF1a, (D1S7W) Rabbit Monoclonal Antibody, CST, Catalog # 99585) staining on these samples overnight at 4°C with a 1:250 dilution, followed by three washes with PBST buffer, and overnight incubation with secondary labelled antibody (Donkey anti-Rabbit IgG conjugated with Alexa Fluor 488, Invitrogen Catalog # A-21206) at 4°C, followed by three washes with PBST buffer. Finally, samples are stained with DAPI at a 1:1000 dilution for one hour, followed by three washes with PBST buffer before imaging. Immunostained samples are stored in PBS with 0.02% sodium azide.

### 4.9. Preparation of bulk RNA-Seq samples

For bulk RNA-Sequencing, we prepare cell-laden 3D collagen matrices in agarose-coated 35 mm plastic dishes. We harvest these by directly adding 1 mL cold Trizol and repeatedly pipetting to completely break up clumps. These samples are first incubated in −80°C for 30 minutes and later thawed on ice. To this, we add 200 μL chloroform, and vortex mix every 30-40 seconds for a total of 10 minutes. During this time, we avoid phase separation. The samples are then centrifuged at 20,000 xg for 40 minutes at 4°C which leads to a clear phase separation. The aqueous phase is collected, to which an equal volume of cold isopropanol and 1 μL glycogen blue is added, vortexed mixed for 10-15 seconds, and finally, incubated on ice for 10 minutes. This mixture is then centrifuged at 20,000 xg for 40 minutes at 4°C. From this, the supernatant is discarded and the RNA fraction is visible as a blue pellet, which is then washed twice in 75% ethanol, with a centrifugation at 14,000 xg for 10 mins at 4°C each time. Finally, the pellets are air-dried at room temperature, and resuspended in 10 μL of nuclease-free water. This represents the pooled bulk RNA fraction. We assess the quality of this using Agilent Bioanalyzer, followed by standard processing with the NEBNext Poly(A) mRNA Magnetic Isolation Module (E7490L) for mRNA isolation and NEBNext® Ultra™ II Directional RNA Library Prep with Sample Purification Beads (E7765L) for polyA-selected RNA library preparation. The samples are sequenced on a NovaSeq 6000 platform using 2×100bp sequencing read length.

### 4.10. Analyses of bulk RNA-Seq Samples

Post sequencing, 18 to 29 million paired-end reads are obtained. FastQC is used to perform the initial quality check. Adapters are trimmed from the reads using cutadapt (-a AGATCGGAAGAGCACACGTCTGAACTCCAGTCA -A AGATCGGAAGAGCGTCGTGTAGGGAAAGAGTGT). The trimmed reads are mapped to the mouse genome (GRCm39) using hisat2 (--rna-strandness R). The mapped reads (91-98% mapping percentage) are counted using featureCounts. DESeq2 is used to perform the read count normalization and differential expression analysis. The plots are generated using R. Genes that show a fold change of 2 or more, with an adjusted p-value of less than 0.05, are considered as differentially expressed for the gene set enrichment analysis. Gene ontology and KEGG pathway enrichment analysis of the differentially expressed genes are done using clusterProfiler.

### 4.11. Preparation of ATAC-Seq samples

For bulk ATAC-Sequencing, we follow a sample preparation protocol similar to that for the bulk RNA sequencing. For harvesting the samples, we apply a collagenase type IV for 30 minutes at 37°C treatment to digest the 3D collagen matrix and retrieve cells. This is centrifuged at 1000 xg for 5 minutes to obtain the cells as a pellet. To this we add a lysis buffer solution comprising of 10 mM Tris pH 7.5m 10 mM NaCl, 4 mM KCl, 0.1% Tween-20, and 0.01% digitonin. The resuspended cell pellet in lysis buffer is continuously tapped for 3 minutes, followed by immediate addition of paraformaldehyde (final conc. 1%), and incubated at room temperature for 5 minutes. To quench the fixation process, 100 mM of glycine buffer is added and incubated for 5 mins. This is followed by a centrifugation at 500 xg for 10 minutes, and resuspension of the subsequently-obtained nuclei pellet in a wash buffer comprising of 10 mM Tris pH 7.5m 10 mM NaCl, 4 mM KCl, and 0.1% Tween-20. We visually confirm the presence of intact nuclei by imaging a 1 μL aliquot from this sample. The remaining sample is pelleted down by a centrifugation at 500 xg for 10 minutes, and entirely used as the input for tagmentation using a transposition reaction mixture that contains 50% of Tagment DNA buffer, 33% of PBS, 0.1% Tween-20, 0.01% digitonin, 5% Tn5 transposase, and the volume being made up with water. This reaction occurs at 37°C for 30 minutes with constant agitation at 1000 rpm. Following this, we commence an overnight reverse-crosslinking reaction at 65°C using a buffer comprising 50 mM Tris pH 8.0, 1 mM EDTA, 0.1% SDS, 0.2 M NaCl, and 2 μl proteinase K in a final volume of 50 μl. Subsequent to this, we follow all remaining steps for library preparation as previously outlined^116^. The sequencing is finally performed on a NovaSeq 6000 platform using 2×100bp sequencing read length.

### 4.12. Analyses of ATAC-Seq Samples

Read quality is assessed using FastQC. Adapters are trimmed using cutadapt (-m 20 -a CTGTCTCTTATACACATCT -A CTGTCTCTTATACACATCT). The trimmed reads are aligned to the mouse genome (GRCm39) using Bowtie2 (--very-sensitive-local). Post-alignment, mitochondrial reads are removed, and PCR duplicates are marked and removed using Picard MarkDuplicates. To account for the Tn5 transposase binding offset, read coordinates are shifted using alignmentSieve (--ATACshift). Accessible chromatin peaks are called using MACS2 (-q 0.05 -f BAMPE --keep-dup all). Peaks overlapping with ENCODE blacklist regions are filtered out using Bedtools. A consensus peak set is generated by merging peaks across samples using Bedops, and reads within these regions are quantified using featureCounts (--fracOverlap 0.2). Differential chromatin accessibility analysis is performed using DESeq2.

### 4.13. Gene Regulatory Network Inference and Differential Network Analysis

A global gene regulatory network (GRN) is constructed by integrating transcriptomic, chromatin accessibility, motif, and TF-activity evidence. GENIE3 is applied to DESeq2-normalized RNA-seq counts to infer TF→target regulatory importance scores. Peak–gene links from ATAC-seq are used to compute RNA–ATAC correlations for each TF–target pair. TF binding evidence is incorporated using motif matches (PWMScan) within accessible regulatory peaks. TF activity is estimated using VIPER, and TF–target correlations from the viper activity matrix are included as an additional regulatory evidence layer. These four components are normalized and combined into a single global regulatory confidence score (final_score), scaled to 0–1. To identify the key drivers of state transitions, condition-specific TF activity is quantified by averaging VIPER scores across replicates and calculating the differential activity for each contrast. A differential GRN is derived by weighting the global final_score by this differential TF activity, yielding an edge delta that represents the condition-dependent gain or loss of regulatory influence. For visualization, TFs are ranked by the magnitude of their differential activity (absolute VIPER score difference). For the top 10 functional drivers in each comparison, the top 100 downstream targets are identified based on the strongest edge deltas. These target lists are subjected to Gene Ontology (GO) enrichment analysis (Biological Process) to determine the dominant function of the regulon. The resulting networks are visualized using ggraph in R, linking each TF to its primary regulated biological pathway rather than individual genes. In these functional GRNs, TF node size is scaled to Total Influence (the cumulative sum of differential edge weights), highlighting regulatory hubs, while node color indicates the direction of activity change (up- or down-regulated).

### 4.14. Global proteome analyses

For global proteome analyses, we follow a sample preparation protocol similar to that for the bulk RNA sequencing but with ten-fold greater cell seeding density, to account for the higher input fraction required for efficient protein extraction. Samples are harvested by digesting the 3D collagen matrix using collagenase type IV for 30 minutes at 37°C, following which cells are retrieved as a pellet by centrifuging at 1000 xg for 5 minutes. The pellet is snap-frozen by a rapid immersion in liquid nitrogen. Protein samples are first reduced with 5 mM TCEP and subsequently alkylated with 50 mM iodoacetamide. The samples are then digested with Trypsin at a 1:50 Trypsin-to-lysate ratio for 16 hours at 37°C. After digestion, the mixture is purified using a C18 silica cartridge and then concentrated by drying in a speed vac. The resulting dried pellet is resuspended in buffer A, which consists of 2% acetonitrile and 0.1% formic acid. For mass spectrometric analysis, all the experiments are performed on an Easy-nLC-1000 system (Thermo Fisher Scientific) coupled with an Orbitrap Exploris 240 mass spectrometer (Thermo Fisher Scientific) and equipped with a nano electrospray ion source. 1μg of peptides sample dissolved in buffer A containing 2% acetonitrile/0.1% formic acid is resolved using Picofrit column (1.8-micron resin, 15cm length). Gradient elution is performed with a 0–38% gradient of buffer B (80% acetonitrile, 0.1% formic acid) at a flow rate of 500nl/min for 96mins, followed by 90% of buffer B for 11 min and finally column equilibration for 3 minutes. Orbitrap Exploris 240 is used to acquire MS spectra under the following conditions: Max IT = 60ms, AGC target = 300%; RF Lens = 70%; R = 60K, mass range = 375−1500. MS2 data are collected using the following conditions: Max IT= 60ms, R= 15K, AGC target 100%. MS/MS data are acquired using a data-dependent top20 method dynamically choosing the most abundant precursor ions from the survey scan, wherein dynamic exclusion is employed for 30s. Samples are processed and RAW files generated are analyzed with Proteome Discoverer (v2.5) against Uniprot reference database. For dual Sequest and Amanda search, the precursor and fragment mass tolerances are set at 10 ppm and 0.02 Da, respectively. The protease used to generate peptides, i.e. enzyme specificity is set for trypsin/P (cleavage at the C terminus of “K/R: unless followed by “P”). Carbamidomethyl on cysteine as fixed modification and oxidation of methionine and N-terminal acetylation are considered as variable modifications for database search. Both peptide spectrum match and protein false discovery rate are set to 0.01 FDR. Differentially abundant proteins (p-value < 0.05) are subjected to functional enrichment analysis to identify over-represented candidates. Protein accession IDs were first mapped to their corresponding gene identifiers. Gene Ontology (GO) enrichment analysis was performed to evaluate enrichment for the Biological Process (BP) GO category. Pathway enrichment analyses are conducted using the Reactome pathway database to identify significantly enriched biological pathways associated with the differentially abundant protein sets. Statistical significance values for enrichment are calculated using a hypergeometric test. Gene ontology groupings and pathways exhibiting p < 0.05 are considered significantly enriched. Enrichment results are represented as chord diagrams representing associations (pathways).

### 4.15. Statistical analyses and sampling quality control

For all statistical comparisons between anoxia and normoxia-exposed cellular populations, we perform an unpaired t-test with appropriate statistical significance values as indicated for each dataset. Specifically for the phase space diagrams, we assign the significance cut-off threshold below a p-value of 0.0001. To determine the appropriate sampling size, we compute the mean, standard deviation, and standard error for sub-groups of different sizes, randomly sampled 100 times from a population of >100 cells for each morphometric parameter across all different 3D collagen matrices, as well as normoxia and anoxia conditions. We learn from this analysis that a sample size of ∼ 60-70 cells appears sufficient to capture the single-cell heterogeneity while dealing with oxo-mechanical phenotypes.

## Acknowledgments

We acknowledge Mridul Gautam, Dr. Jasmine Dhall, Dr. Vinay Dubey, Sarayu Beri, Uttkarsh Ayyangar, Astha Mukhopadhyay, Nivedita Chaudhary, and Atriya Mazumdar for their helpful assistance with the experiments and sample preparation protocols. We are grateful to Prof. Satyajit Mayor and Dr. Ramray Bhat for their numerous critical inputs and advice over the course of this study. We also acknowledge the valuable discussions with Dr. Sudarshan Gadadhar, Prof. Madan Rao, Prof. Thomas Angelini, Prof. Sujit Datta, Prof. Srikala Raghavan, Prof. L S Shashidhara, and Prof. Upinder Bhalla. We thank Dr. Sudarshan Gadadhar for generously sharing the NIH-3T3 cells. We thank the Central Imaging and Flow Cytometry Facility (CIFF) for access to confocal microscopy. We thank Dr. Awadesh Pandit, Lakshminarayanan CP, and the Next Generation Sequencing Facility staff at NCBS. A set of schematic figures were prepared using BioRender. TB acknowledges an intramural research grant from NCBS and research funding from Anusandhan National Research Foundation, Govt. of India (CRG/2021/008869). This work was supported by Nikon India. MS acknowledges personal support from TIFR graduate school.

## Author Contributions

TB, DP, and MS conceptualized the study. MS developed the methodology with guidance from TB. MS performed all experiments and analyzed the data. NH performed the multi-omics analyses with guidance from DP. MS validated the findings and curated the final data. MS prepared the figures. MS wrote the manuscript with feedback from TB and DP. TB and DP secured primary funding, contributed resources, and arranged infrastructural support. TB supervised the overall study.

## Data and materials availability

All data are available in the main text or the supplementary materials. The RNA-Seq and ATAC-Seq data corresponding to this study are deposited to the NCBI SRA database, with the BioProject ID PRJNA1430130. The proteomics data corresponding to this study are deposited to the ProteomeXchange Consortium via the PRIDE partner repository with the dataset identifier PXD075045.

## Competing interests

Authors declare that they have no competing interests.

## Supplementary Information

**SI Fig 1.**
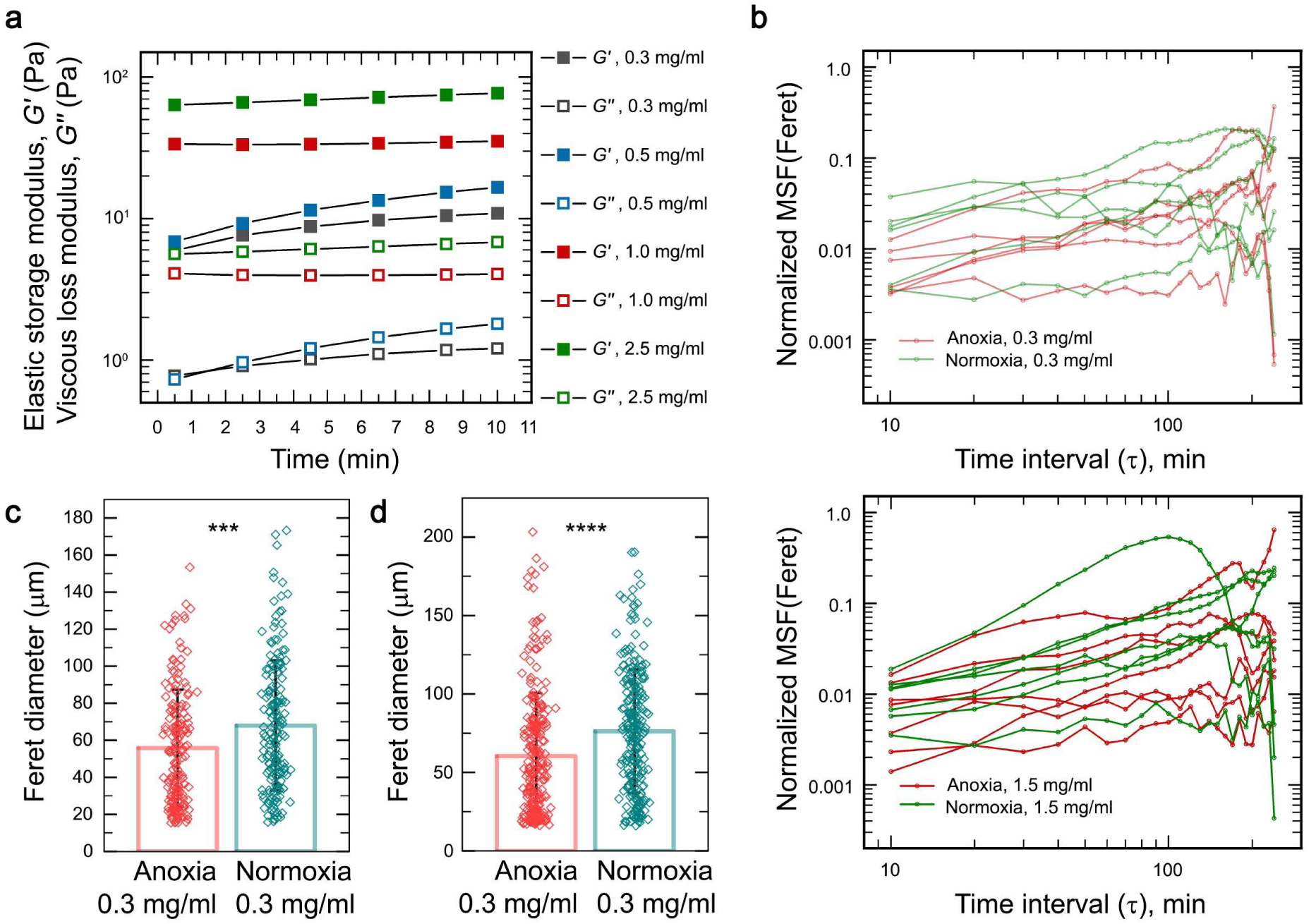
(**a**) Shear rheology reveals the tunable, concentration-dependent viscoelastic properties of 3D collagen hydrogels. G’ represents the elastic storage modulus (elastic solid-like behavior), whereas G’’ represents the viscous loss modulus (viscous fluid-like behavior). (**b**) Quantification of single cell morphology (feret diameter) from timelapse imaging of cells embedded in either low (0.3 mg/ml) or high (1.5 mg/ml) fiber density collagen matrices, maintained under either anoxia or normoxia, following an initial 12 hrs of incubation under the respective conditions. Each time trace represents an individual cell. n >= 6 individual cells for all conditions, imaged using time lapse microscopy for = 4 hours each. Data represented as a mean squared fractional change (MSF) in feret diameter between consecutive time frames, normalized to the feret diameter in the first frame of reference. (**c** and **d**) Single cell feret diameter measurements for cells embedded in 0.3 mg/ml collagen fiber density matrices, maintained under either anoxia or normoxia, performed at (**c**) 40 hours post-seeding (n > 170 individual cells for each condition, data represented as mean +/− s.d., statistical significance calculated using an unpaired t-test, with p-value < 0.001) and (**d**) 10-fold higher cell density (n > 200 individual cells for each condition, data represented as mean +/− s.d., statistical significance calculated using an unpaired t-test, with p-value < 0.0001), both of which also show a difference between the population-level cell morphologies across anoxia and normoxia.

**SI Fig. 2.**
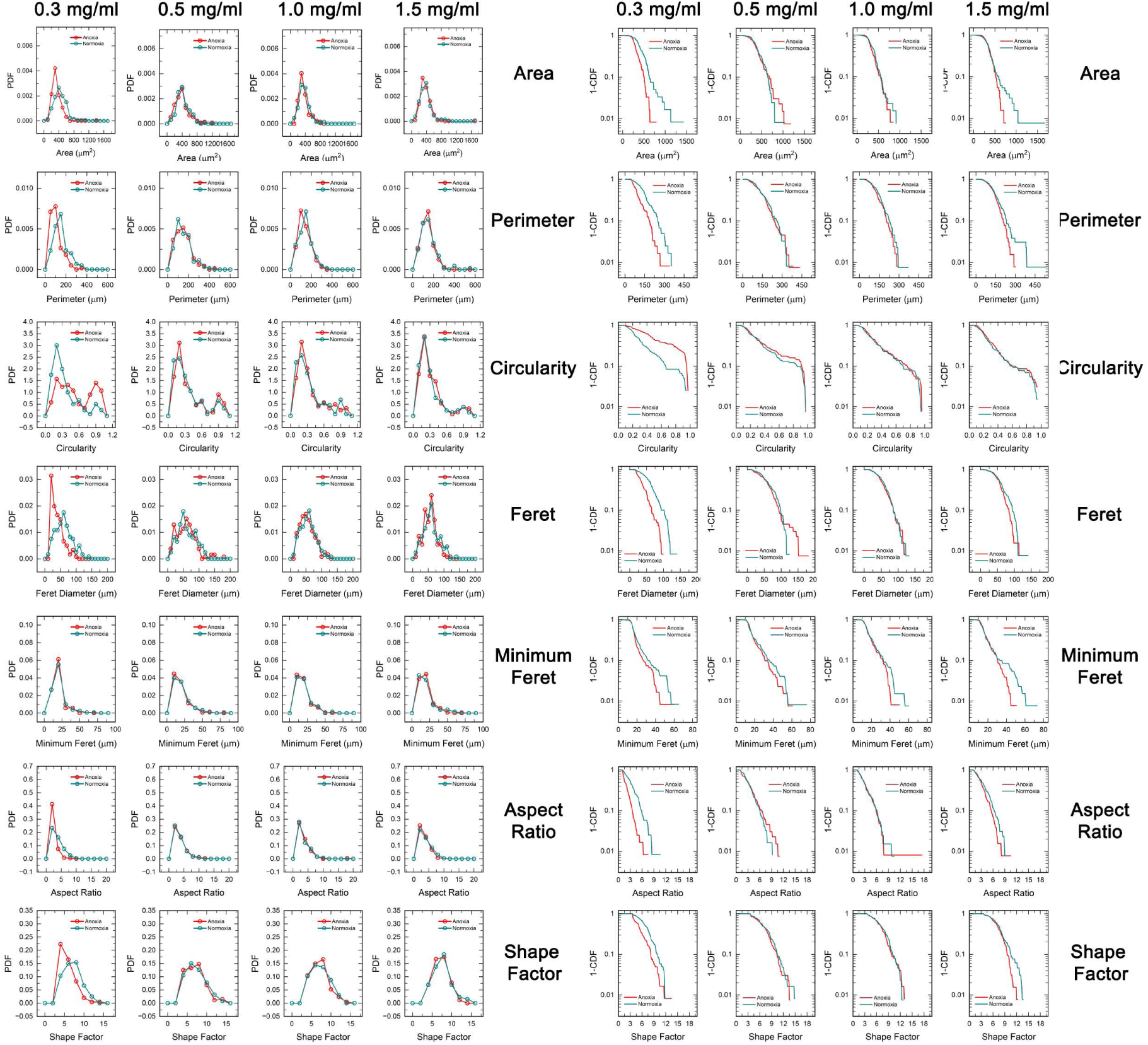
Population-level analyses of morphometrics performed on cells across various oxo-mechanical regimes, represented as probability density functions and complementary cumulative distribution functions. n >= 120 individual cells for each condition.

**SI Fig. 3.**
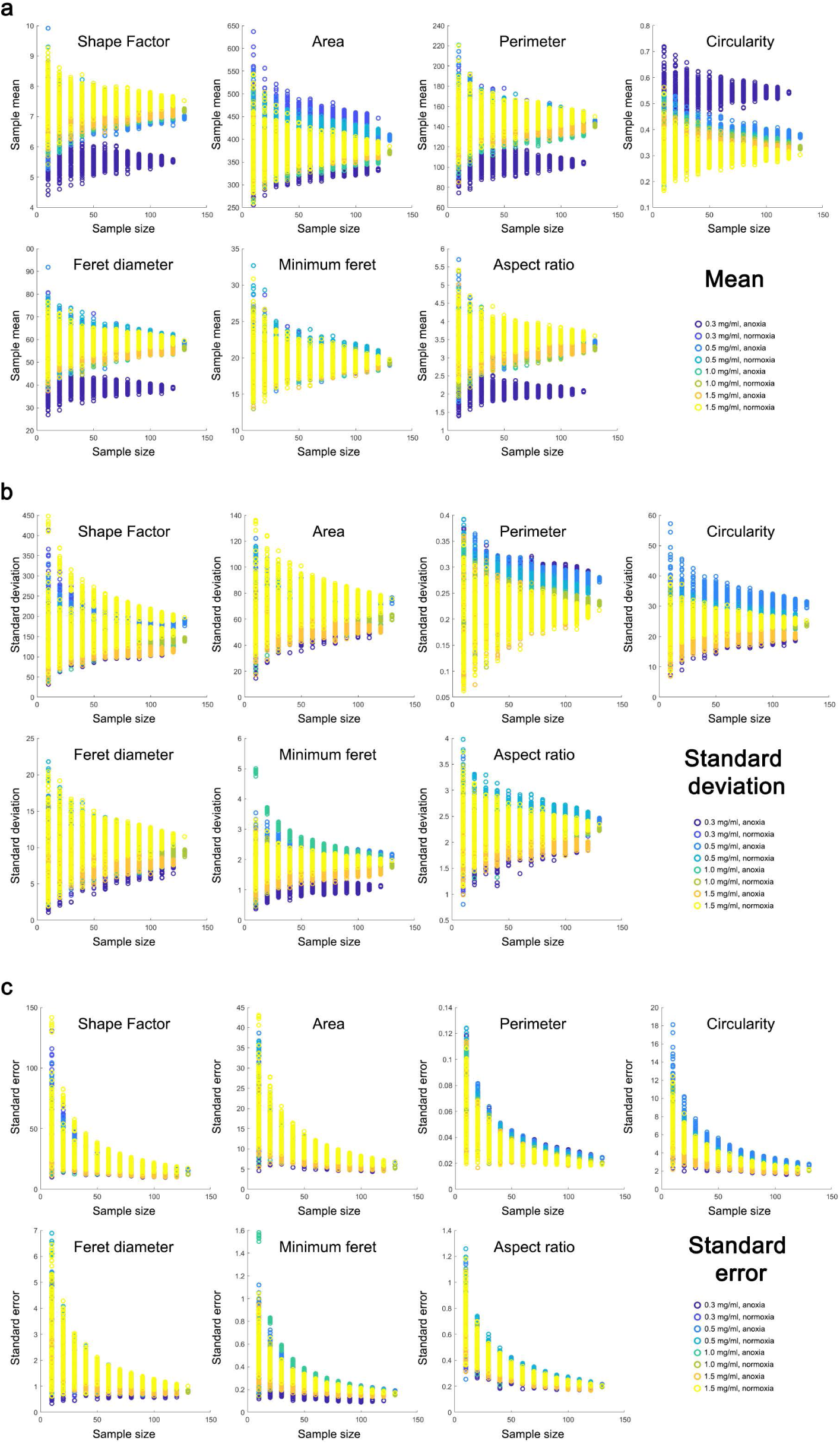
Sampling robustness tests performed on single cell morphometric data obtained from various oxo-mechanical regimes, shown here by assessing the (**a**) population average (mean), (**b**) standard deviation, and (**c**) standard error, using varying sample sizes randomly picked from the overall population, for which all three parameters are calculated, with the process being iterated 100 times for each such sample size. n >= 120 individual cells for each condition.

**SI Fig. 4.**
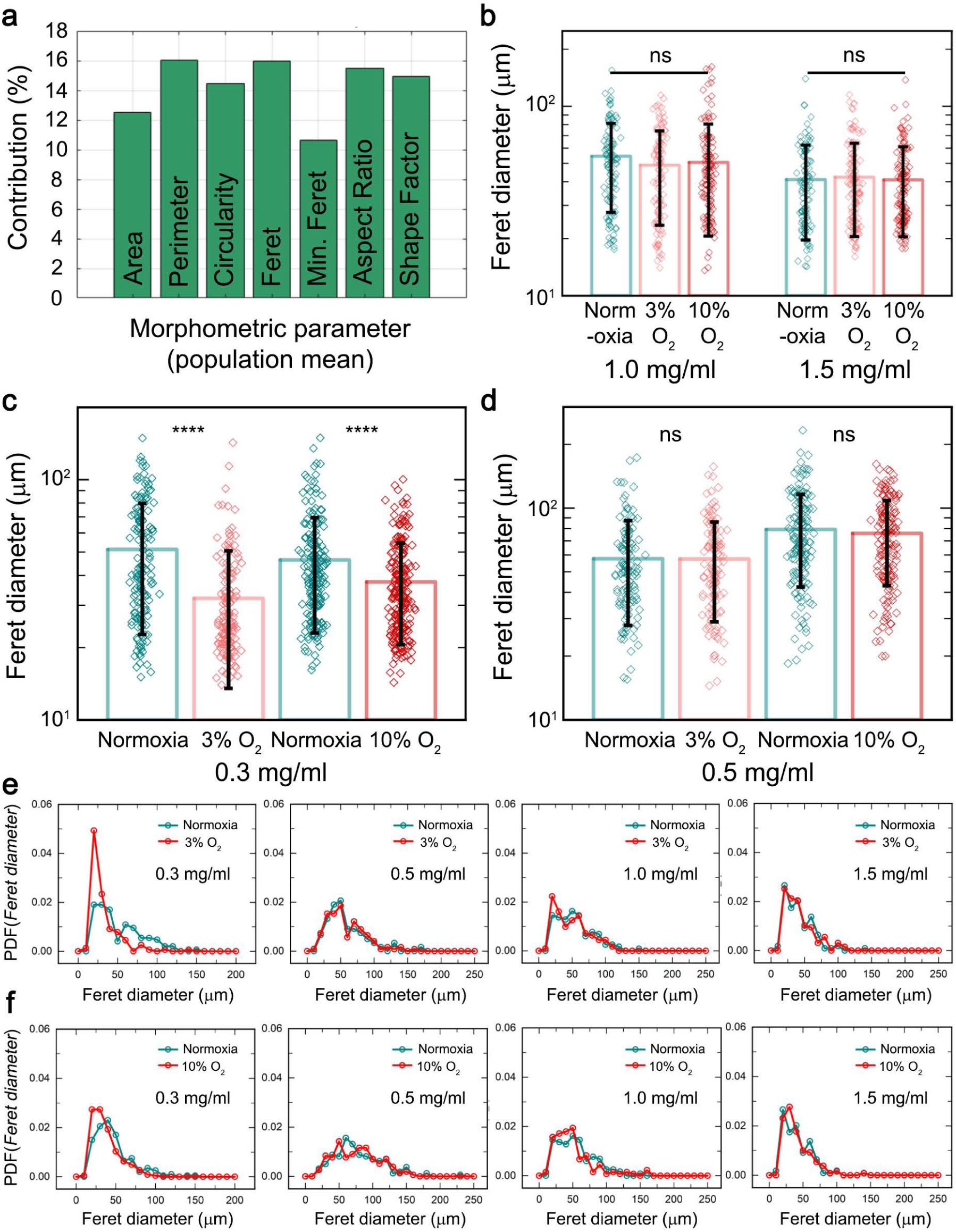
(**a**) Relative contribution of each morphometric parameter towards the principal component analysis’ PC1. n >= 120 individual cells for each condition. (**b-d**) Individual cell-level data from cells embedded within varying collagen fiber density matrices, and maintained under either 3% or 10% oxygen partial pressures, shown alongside their normoxia counterparts – reflecting a clear shift in the cellular morphology from more rounded-up under hypoxia to more elongated under normoxia. (**e** and **f**) Population-level analyses of feret diameters, represented as a probability density function, obtained from cells embedded in different collagen fiber density matrices, maintained under either (**e**) 3% or (**f**) 10% oxygen partial pressures, with a normoxia control. n > 100 individual cells for each condition.

**SI Fig. 5.**
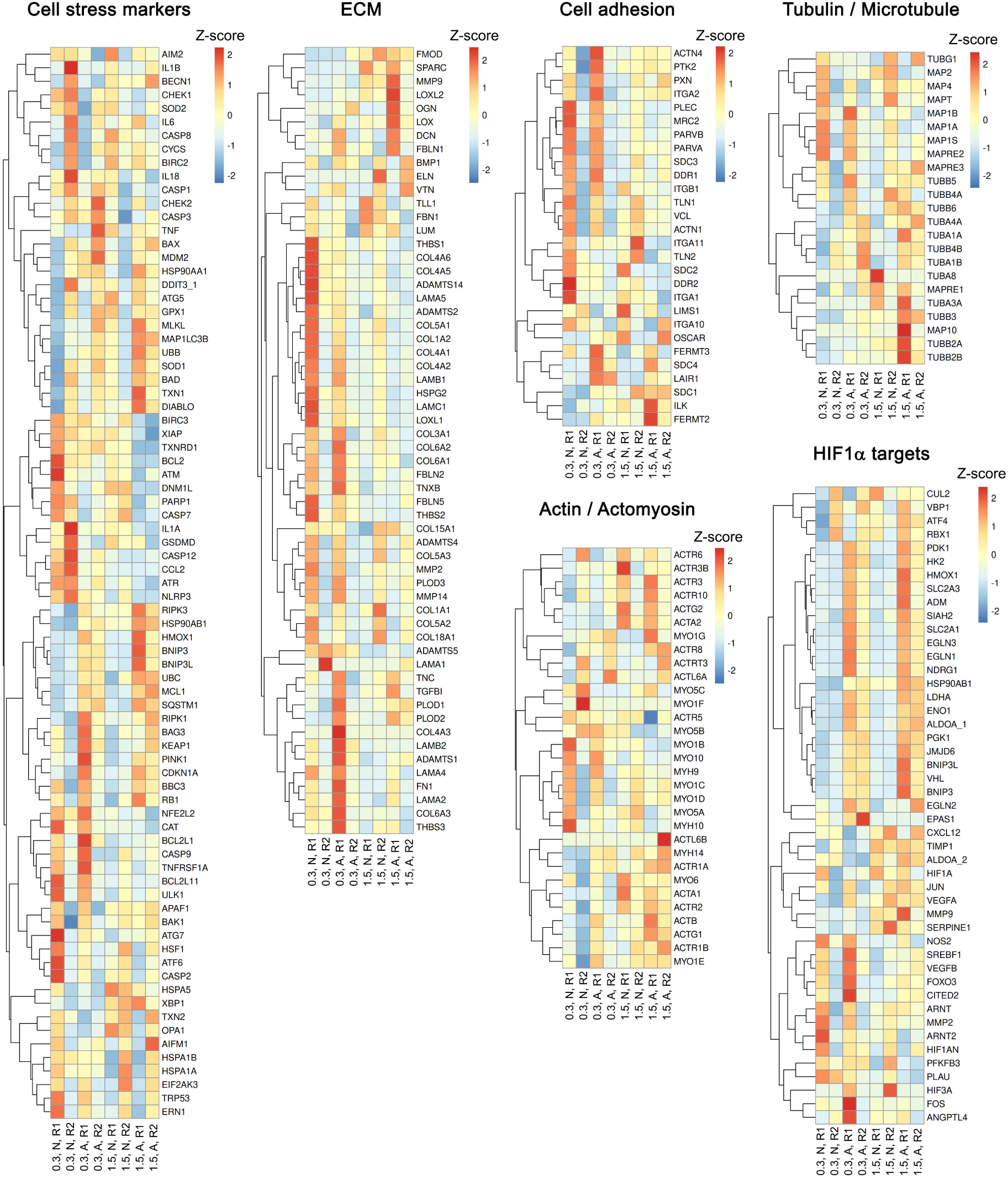
Heatmaps representing z-scores calculated using relative gene expression levels for different biological functionalities based on the transcriptome analyses performed on cells embedded in either poorly-reinforced (0.3 mg/ml) or well-reinforced (1.5 mg/ml) mechanical milieus, maintained under either anoxia or normoxia.

**SI Fig. 6.**
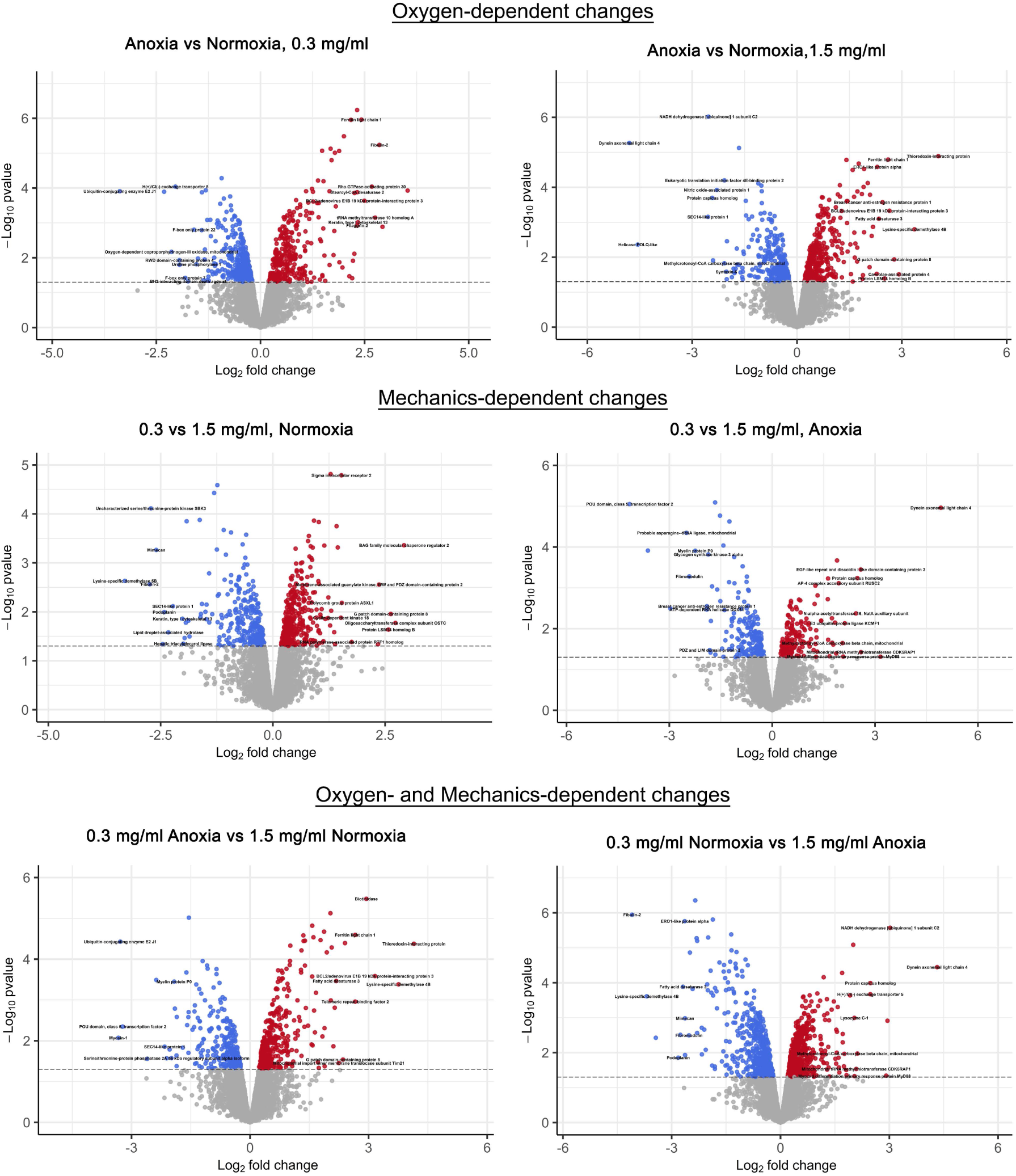
Volcano plots representing differentially-abundant proteins based on the quantitative proteomics analyses performed on cells embedded in either poorly-reinforced (0.3 mg/ml) or well-reinforced (1.5 mg/ml) mechanical milieus, maintained under either anoxia or normoxia. Statistical threshold for identifying differentially-abundant proteins across all pairwise comparisons was set at a p-value < 0.05.

**SI Fig. 7.**
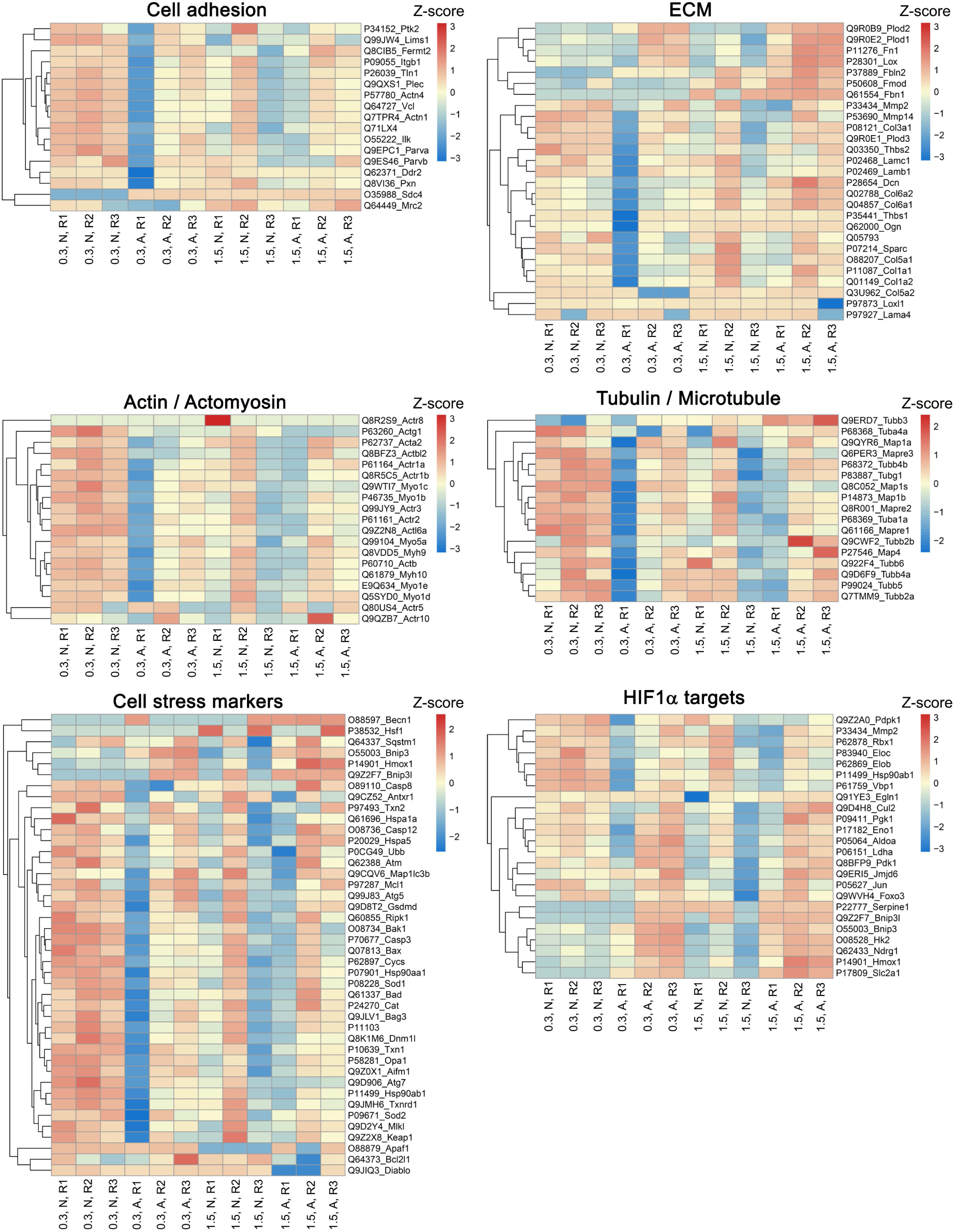
Heatmaps representing z-scores calculated using relative protein abundance levels for different biological functionalities based on the proteomics analyses performed on cells embedded in either poorly-reinforced (0.3 mg/ml) or well-reinforced (1.5 mg/ml) mechanical milieus, maintained under either anoxia or normoxia.

**SI Fig. 8.**
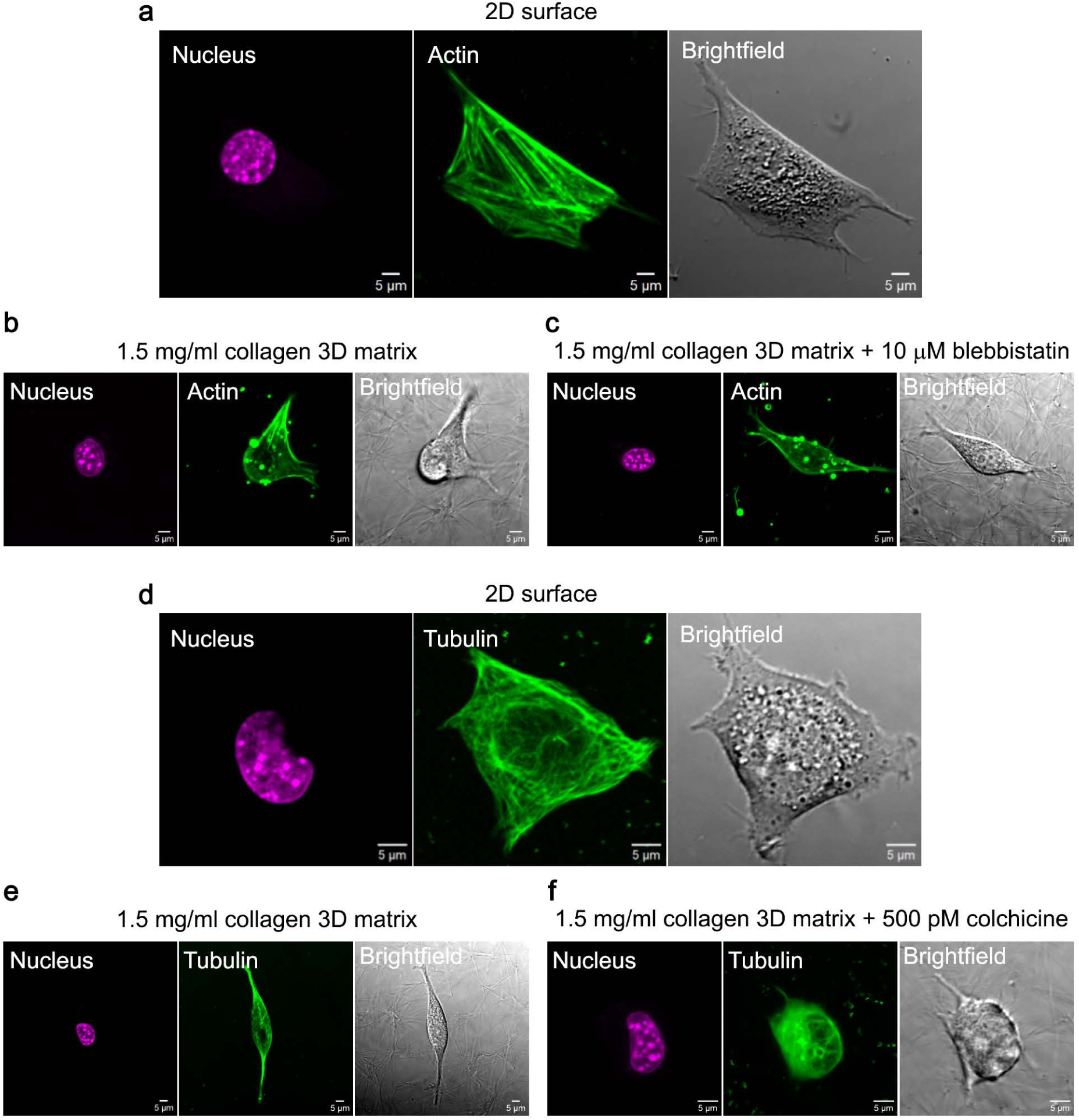
Cells live-stained for (**a-c**) actin and (**d-f**) microtubules. While cells adherent on 2D glass surfaces exhibit (**a**) prominent actin stress fibers and (**d**) well-resolved, densely-packed microtubule networks, cells adherent in 3D collagen gels largely exhibit (**b**) cortical actin and (**e**) sparser microtubular networks. (**c**) Disruption of the actomyosin contractility using blebbistatin does not significantly alter the cortical actin arrangements, whereas, (**f**) disruption of microtubular stability using colchicine manifests as poorly-defined, diffuse tubular structures.

**SI Fig. 9.**
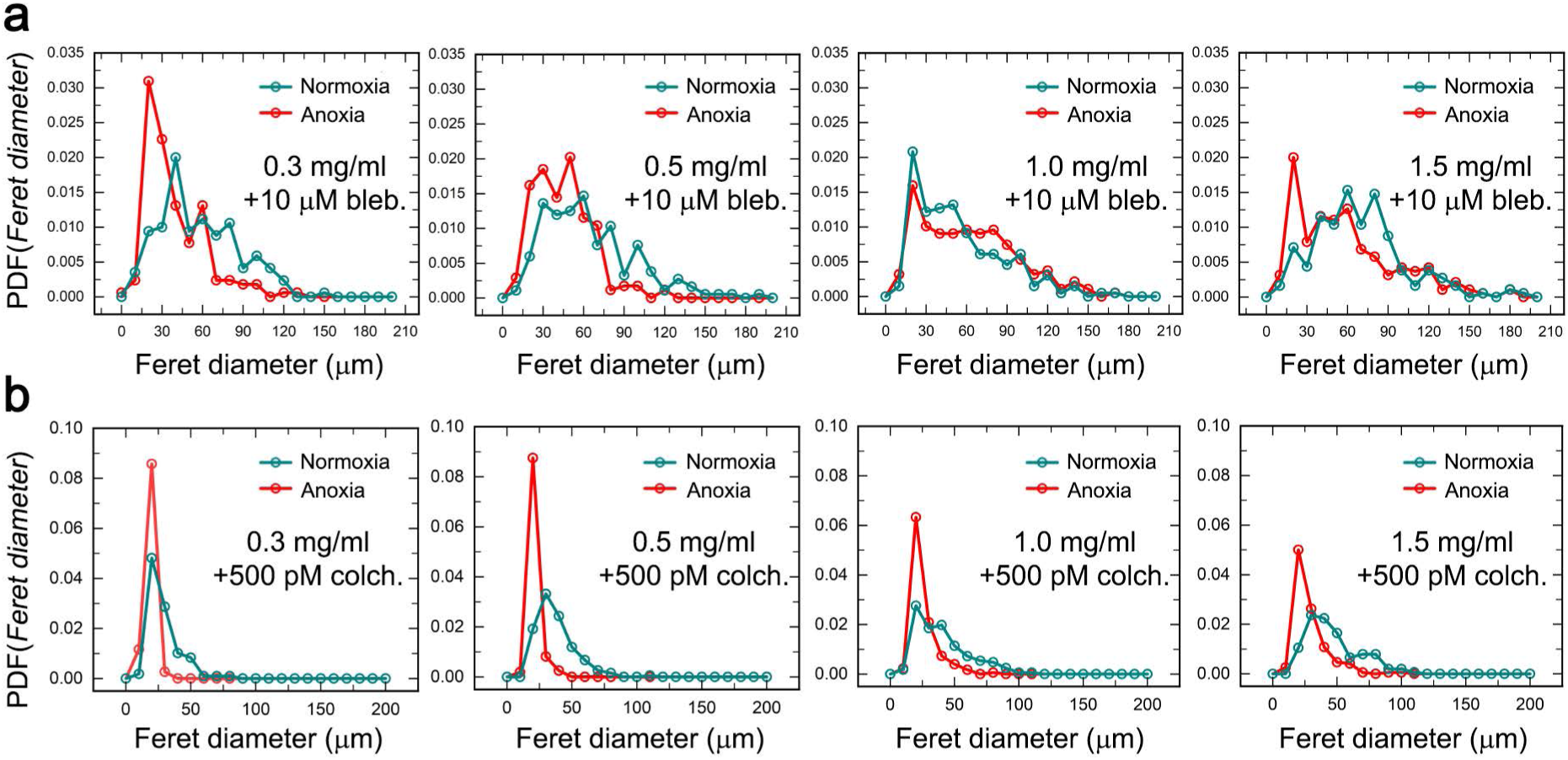
Population-level analyses of feret diameters, represented as probability density function, obtained from cells embedded in different collagen fiber density matrices and maintained under either anoxia or normoxia, treated with either (**a**) 10 μM blebbistatin to disrupt actomyosin contractility (n > 150 individual cells for each condition) or (**b**) 500 pM colchicine to disrupt microtubule stability (n > 100 individual cells for each condition).

**SI Fig. 10.**
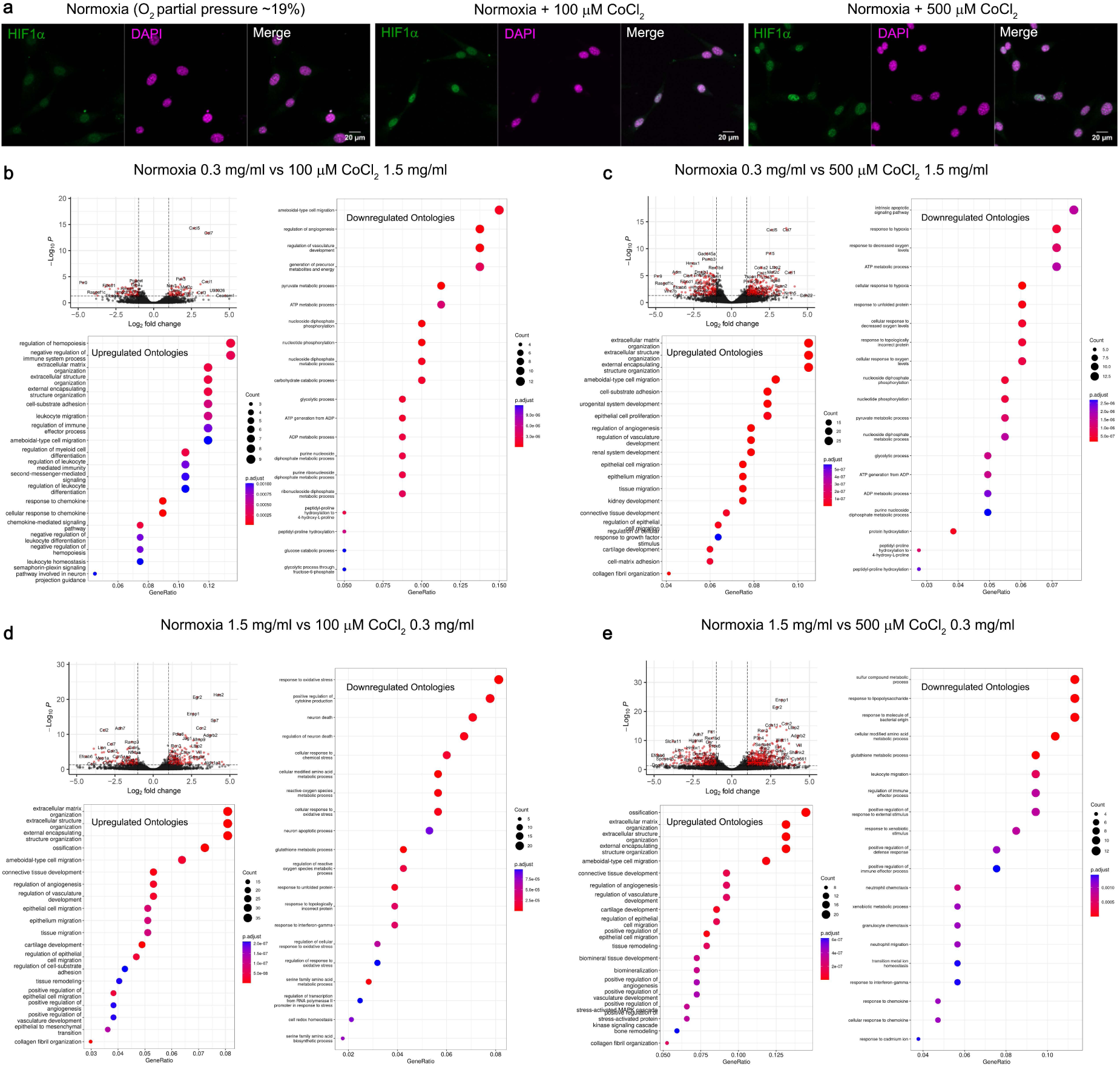
(**a**) Immunostaining for HIF1α reveals that cobalt chloride treatments enhance nuclear localization of HIF1α even under normoxic conditions, which is indicative of cellular hypoxic signaling responses. (**b**-**e**) Volcano plots and ontological associations with biological processes for differentially-expressed genes, based on transcriptome analyses performed on cells embedded in either poorly-reinforced (0.3 mg/ml) or well-reinforced (1.5 mg/ml) mechanical milieus, treated with either 100 μM or 500 μM of the chemical hypoxia mimic cobalt chloride. Statistical threshold for identifying differentially-expressed genes across all pairwise comparisons was set as |log2FC| >= 1, i.e., 2-fold change in expression levels, with adjusted p-value < 0.05.

**SI Fig. 11.**
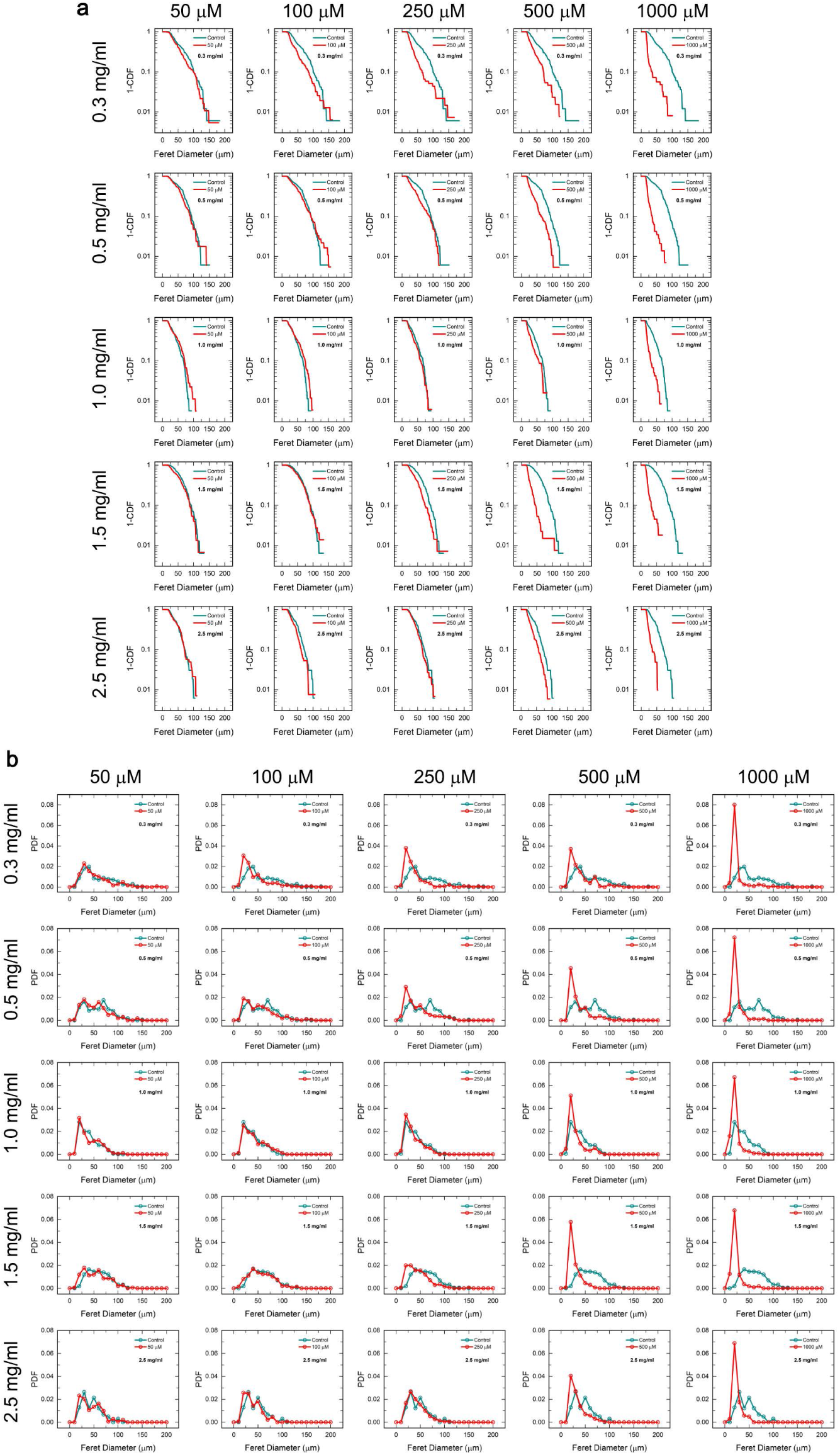
Population-level analyses of cellular morphology using complementary cumulative distribution functions for cells across five different collagen fiber density matrices while being subjected to five different grades of chemically-induced hypoxia, showing how elevated hypoxic signaling pushes the population towards a more rounded-up morphology, whereas, increased mechanical reinforcement in the form of high collagen fiber density counteracts this effect - albeit, within limits. Analyses represented as (**a**) complementary cumulative distribution functions and (**b**) probability density functions. n > 100 individual cells for each condition.

**SI Fig. 12.**
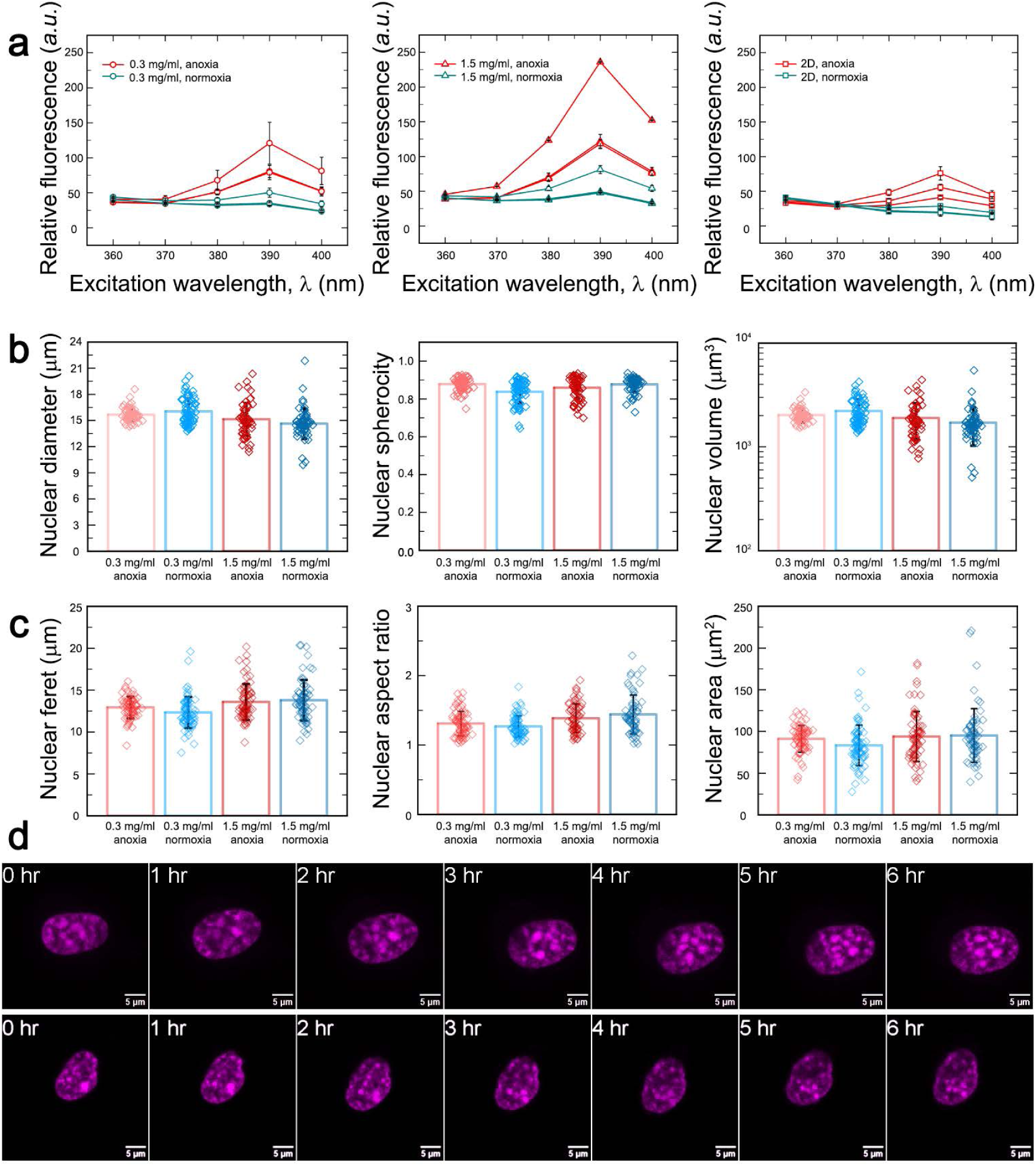
(**a**) Time-resolved fluorescence measurements of intracellular oxygen concentrations, performed on cells embedded in 0.3 mg/ml 3D collagen matrix, 1.5 mg/ml 3D collagen matrix, and plated on a 2D collagen-coated glass surface, all of which are maintained under either anoxia or normoxia, as specified. n = 3 independent replicates, data represented as mean +/− s.d. (**b-c**) Quantitative analyses of nuclear morphology, performed using either (**b**) 3D segmentation or (**c**) 2D maximum intensity projections, showing no significant differences across different oxo-mechanical regimes. n > 55 individual cells for each condition. (**d**) Live imaging of Hoechst-stained nuclei from cells embedded in 1.5 mg/ml collagen matrices under normoxia.

**SI Fig. 13.**
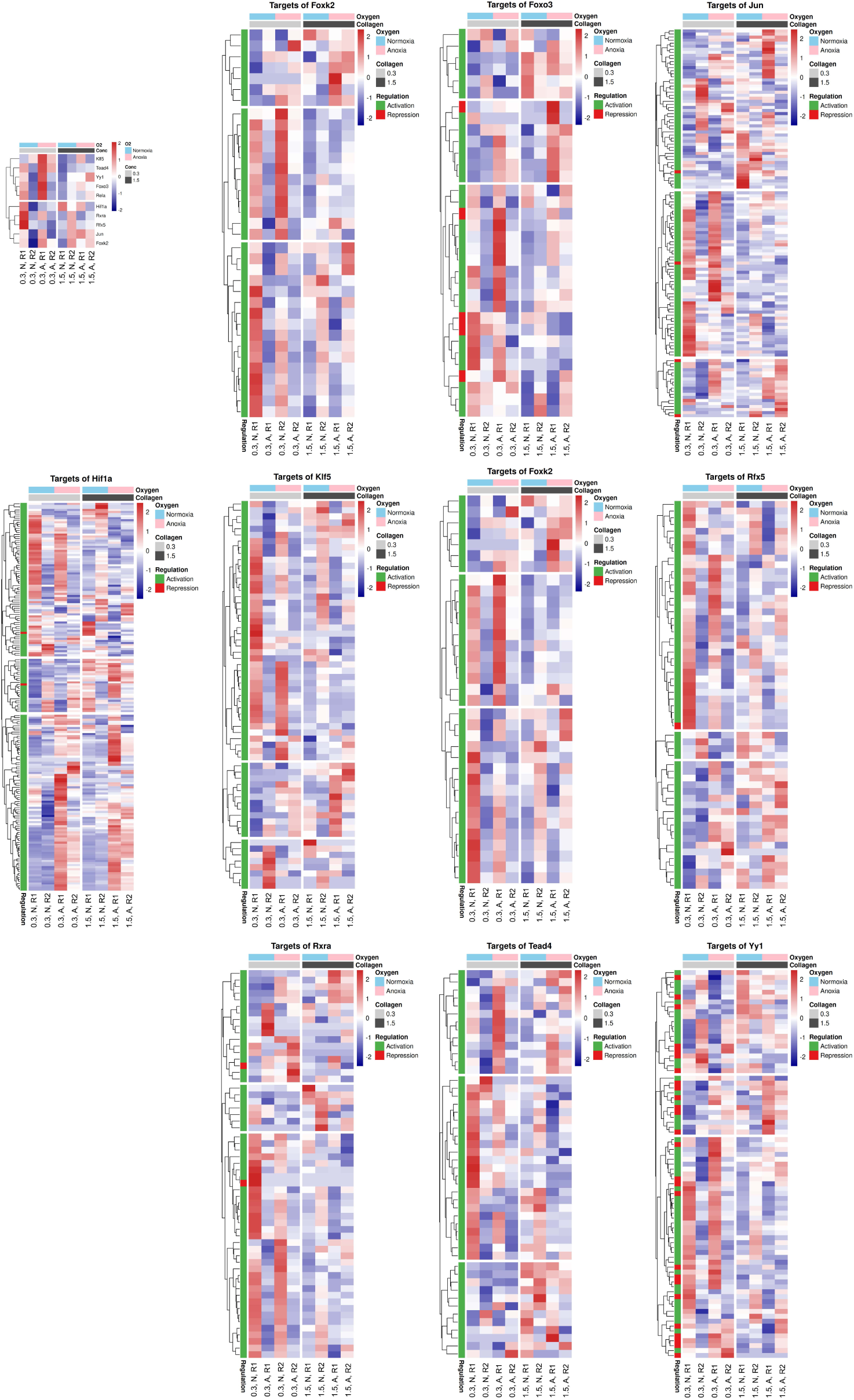
Activity profiles of key transcription factors based on the gene regulatory network analyses, indicating their potential roles in either activating or repressing the expression of their canonically-associated targets across different oxo-mechanical regimes.

**SI Fig. 14.**
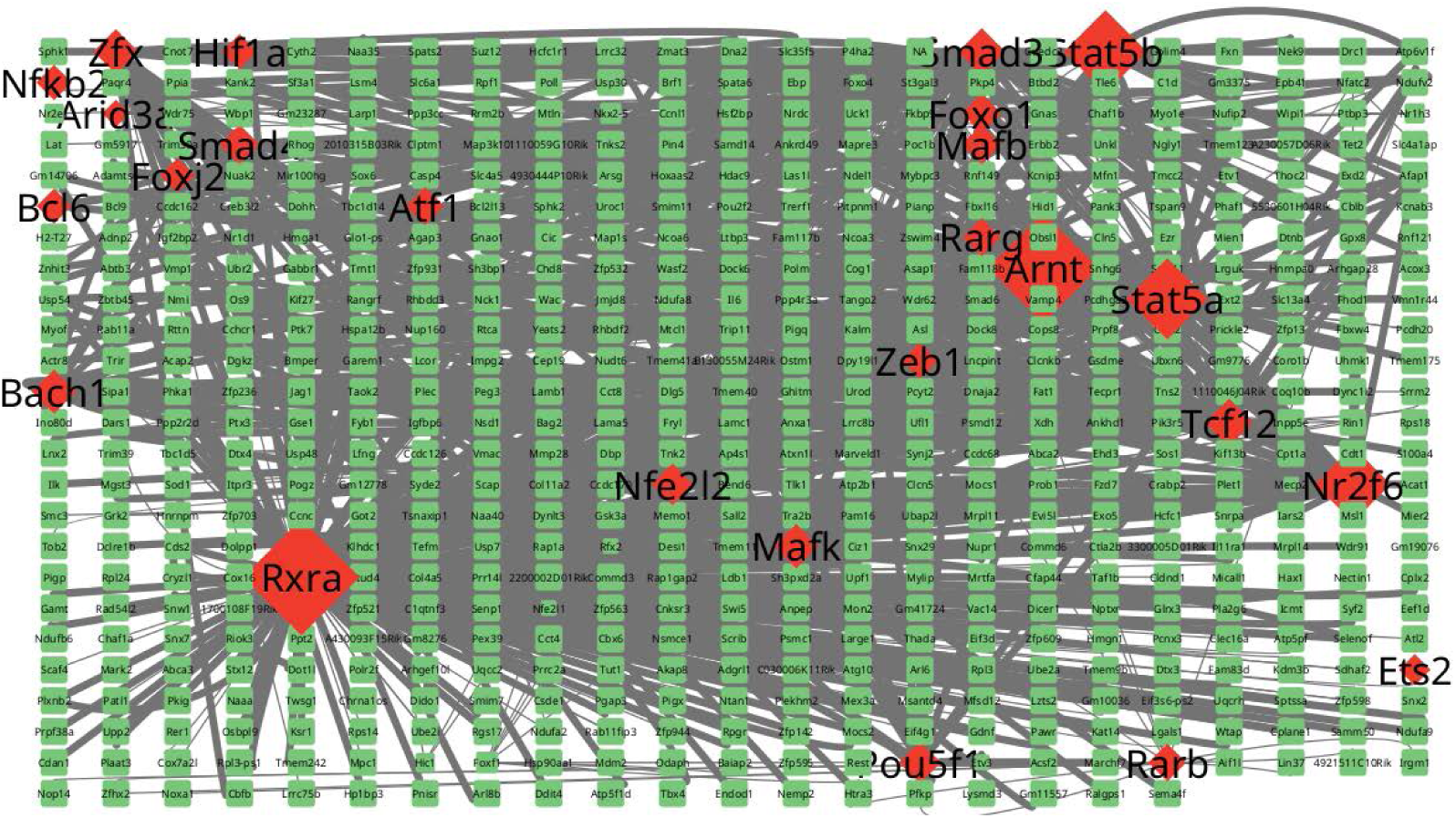
A global GRN spanning all four oxo-mechanical regimes (0.3 mg/ml anoxia, 0.3 mg/ml normoxia, 1.5 mg/ml anoxia, and 1.5 mg/ml normoxia).

## Description of SI Tables

**SI Table 1**: Summary of transcription factors (TFs) ranking across all oxo-mechanical regimes.

**SI Table 2**: Rankings of TFs targets for the pairwise comparison – 1.5 mg/ml normoxia vs 0.3 mg/ml normoxia

**SI Table 3**: Rankings of TFs targets for the pairwise comparison – 1.5 mg/ml anoxia vs 0.3 mg/ml anoxia

**SI Table 4**: Rankings of TFs targets for the pairwise comparison – 0.3 mg/ml anoxia vs 0.3 mg/ml normoxia

**SI Table 5**: Rankings of TFs targets for the pairwise comparison – 1.5 mg/ml anoxia vs 1.5 mg/ml normoxia

**SI Table 6**: Rankings of TFs targets for the pairwise comparison – 0.3 mg/ml anoxia vs 1.5 mg/ml normoxia

**SI Table 7**: Rankings of TFs targets for the pairwise comparison – 0.3 mg/ml normoxia vs 1.5 mg/ml anoxia

**SI Table 8**: Gene ontology classification for biological processes based on the TFs targets for the pairwise comparison – 1.5 mg/ml normoxia vs 0.3 mg/ml normoxia

**SI Table 9**: Gene ontology classification for biological processes based on the TFs targets for the pairwise comparison – 1.5 mg/ml anoxia vs 0.3 mg/ml anoxia

**SI Table 10**: Gene ontology classification for biological processes based on the TFs targets for the pairwise comparison – 0.3 mg/ml anoxia vs 0.3 mg/ml normoxia

**SI Table 11**: Gene ontology classification for biological processes based on the TFs targets for the pairwise comparison – 1.5 mg/ml anoxia vs 1.5 mg/ml normoxia

**SI Table 12**: Gene ontology classification for biological processes based on the TFs targets for the pairwise comparison – 0.3 mg/ml anoxia vs 1.5 mg/ml normoxia

**SI Table 13**: Gene ontology classification for biological processes based on the TFs targets for the pairwise comparison – 0.3 mg/ml normoxia vs 1.5 mg/ml anoxia

**SI Table 14**: Summary of the top 1000 nodes identified in the gene regulatory network (GRN) analyses.

## Description of SI Videos

**SI Video 1**: Timelapse imaging of a cell embedded in a 0.3 mg/ml collagen matrix and incubated under anoxia. Collagen fibers are visualized using reflectance microscopy, while the cell body has been stained using the viability-indicating dye Calcein Green-AM.

**SI Video 2**: Timelapse imaging of a cell embedded in a 0.3 mg/ml collagen matrix and incubated under normoxia. Collagen fibers are visualized using reflectance microscopy, while the cell body has been stained using the viability-indicating dye Calcein Green-AM.

**SI Video 3**: Timelapse imaging of a cell embedded in a 1.5 mg/ml collagen matrix and incubated under anoxia. Collagen fibers are visualized using reflectance microscopy, while the cell body has been stained using the viability-indicating dye Calcein Green-AM.

**SI Video 4**: Timelapse imaging of a cell embedded in a 1.5 mg/ml collagen matrix and incubated under normoxia. Collagen fibers are visualized using reflectance microscopy, while the cell body has been stained using the viability-indicating dye Calcein Green-AM.

**SI Video 5**: Timelapse imaging of nuclei stained with Hoechst from cells embedded in a 1.5 mg/ml collagen matrix and incubated under normoxia.

## Notes

### Competing Interest Statement

The authors have declared no competing interest.

